# Toxin-mediated depletion of nicotinamide dinucleotides drives persister formation in a human pathogen

**DOI:** 10.1101/2023.09.28.559889

**Authors:** Isabella Santi, Raphael Dias Teixeira, Pablo Manfredi, Daniel Spiess, Guillaume Mas, Alexander Klotz, Nicola Zamboni, Sebastian Hiller, Urs Jenal

## Abstract

Toxin-antitoxin (TA) systems are widespread in bacteria and are implicated in genome stability, virulence, phage defense and persistence. Although TA systems encompass a large variety of molecular activities and cellular targets, their physiological role and regulatory mechanisms are often unclear^1,2^. Here, we show that a RES domain TA system increases the survival of the human pathogen *P. aeruginosa* during antibiotic treatment by generating a subpopulation of highly drug-tolerant persisters. The NatT toxin is an NAD phosphorylase, which leads to strong depletion of NAD and NADP in a subpopulation of cells. Actively growing *P. aeruginosa* cells effectively compensate for toxin-mediated NAD deficiency by inducing the NAD salvage pathway. In contrast, under nutrient-limited conditions, NatT generates NAD-depleted cells that give rise to drug tolerant persisters during outgrowth. Structural and biochemical analyses of active and inactive NatR-NatT complexes reveal how changes in NatR-NatT interaction controls toxin activity and autoregulation. Finally, we show that the NAD precursor nicotinamide blocks NatT activity and eliminates persister formation, exposing powerful metabolic feedback control of toxin activity. The findings that patient isolates contain *natT* gain-of-function alleles and that NatT increases *P. aeruginosa* virulence, argue that NatT contributes to *P. aeruginosa* fitness during infections. These studies provide mechanistic insight into how a TA system promotes pathogen persistence by disrupting essential metabolic pathways during nutrient stress.

## Introduction

Antibiotic tolerance is being recognized as an important survival strategy that bacterial pathogens use to escape the lethal effects of bactericidal antibiotics^3–5^. Subpopulations of drug tolerant bacteria, called persisters, not only promote pathogen survival and recurrent infections, but they also provide protected reservoirs for the development of antibiotic resistance^5–7^. Although the molecular basis of persisters is still poorly understood, it is assumed that they can originate from small subpopulations of metabolically dormant cells in response to nutrient exhaustion or to other forms of stress^4,8^. Generally, persisters account for only a small fraction of bacterial populations, making mechanistic studies of drug tolerance challenging. However, hyper-persister lineages evolve during chronic infections^5,9,10^ or in laboratory populations repeatedly exposed to antibiotics^3,5,11^, offering entry points into this phenomenon by providing access to larger persister populations and by exposing potential drivers of drug tolerance.

We have recently isolated hyper-persister variants of *P. aeruginosa*, an important human pathogen causing acute and chronic infections^5^. One of the mutations conferring drug tolerance was mapped to *PA1030*, a gene encoding a RES domain containing toxin that is part of a type II toxin-antitoxin (TA) module^1^. Its upstream neighbor, *PA1029*, encodes a small putative transcription factor with an N-terminal HTH domain and a C-terminal Xre/MbcA/ParS-like, toxin binding domain (**Fig. 1a**). Based on their biological function outlined below, we renamed these genes *natR* (<u>NA</u>D^+^ degrading toxin <u>r</u>epressor) and *natT* (<u>NA</u>D^+^ degrading toxin). RES domain proteins (COG5654) are widespread in bacteria, including important human pathogens like *Mycobacterium tuberculosis*, *Yersinia pestis*, *Brucella abortus*, *Legionella pneumophila*, *Bordetella pertussis* or *Burkholderia pseudomallei*^12–14^. Importantly, RES toxins were shown to cleave nicotinamide dinucleotide (NAD) or to use NAD to modify targets through an ADPribosylation reaction, indicating that RES-based TA systems act by limiting central nicotina-mide-based cofactors or by allosterically modifying downstream targets^12,14^. Recent studies have also implicated NADase toxins in bacterial defense against bacteriophages through a self-destructive process called abortive infection^15–18^. However, known phage defense systems targeting NAD either engage TIR (Toll/IL-1 receptor) or SIR2 (sirtuin) NADases, ancient immune modules that have spread from bacteria to animals and plants, where they have adopted important roles in innate immune defenses^19^.

**Figure 1.**
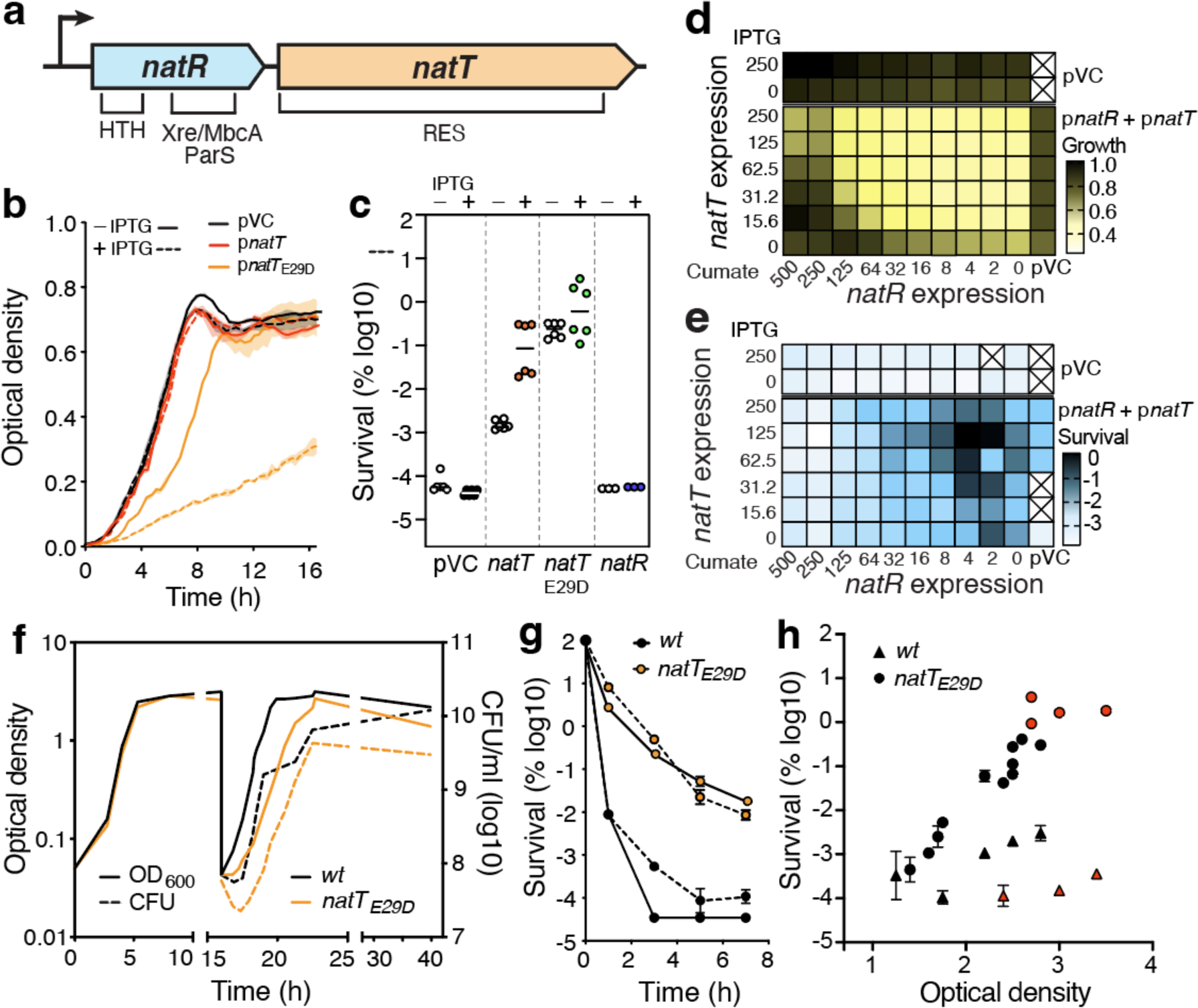
The NatT toxin confers drug tolerance to *P. aeruginosa*. **a)** Schematic of the *natR-natT* locus and domain structure of the NatT toxin and NatR antitoxin. **b)** Expression of *natT* impairs *P. aeruginosa* growth. Cultures of a Δ*natT* mutant containing plasmids with IPTG-inducible *natT* alleles were grown in LB medium with or without IPTG (average ± SEM, n>3) (pVC=control plasmid). **c)** Expression of *natT* increases antibiotic tolerance. Cultures of *P. aeruginosa* containing plasmids with IPTG-inducible *natT* or *natR* alleles were grown in LB medium without or with IPTG. Fractions of surviving cells were determined after three hours of treatment with tobramycin (16 μg/ml). Median values are indicted (n≥3) (pVC=control plasmid). **d,e)** Expression of *natT* limits *P. aeruginosa* growth and increases tolerance. A Δ*natR-*Δ*natT* mutant carrying plasmids with an IPTG-inducible copy of *natT* and a cumate-inducible copy of *natR* was grown in LB supplemented with different concentrations of IPTG and cumate, as indicated. Growth (**d**) and tolerance to ciprofloxacin (2.5 μg/ml) (**e**) were recorded for each combination. Heat maps in (**d**) show ODs reached in the presence of the inducers normalized to the highest OD value of the analyzed group. Heat maps in (**e**) show the fraction of surviving cells after treatment with ciprofloxacin (pVC=control plasmid). **f**) Growth of *P. aeruginosa* wild type and *natTE29D* mutant in LB complex media. Stationary phase cultures were diluted into fresh medium at 15 hrs post inoculation. **g)** Survival of *P. aeruginosa* wild type and *natTE29D* mutant upon exposure to tobramycin (16 μg/ml, solid lines) and ciprofloxacin (2.5 μg/ml, stippled lines) (average ± SEM, n=3). **h)** Tolerance is growth phase-dependent. Cultures of *P. aeruginosa* wild type (triangles) and *natTE29D* mutant (circles) were harvested at different stages of growth (indicated by OD) and exposed to tobramycin (16 μg/ml) for 3 hrs. Fractions of surviving cells from stationary phase are indicated in red (average ± SEM, n=3).

Most RES domains are part of bacterial TA systems^13^. Although TA systems have been shown to play prominent roles in abortive infection and phage defense, none of the RES-domain TA systems has been implicated in bacterial immunity, leaving the physiological role of this large protein family unresolved. Some members have recently been implicated in solvent tolerance, plasmid stability or stress-mediated growth inhibition^12,14,20–22^. *E.g.*, the MbcA-MbcT system of *M. tuberculosis* is significantly upregulated in a variety of stress conditions, including persister cells^23^, hypoxic stress^24^, starvation^25^, and in human macrophages^26,27^. Similarly, the *natR-natT* module of *P. aeruginosa* is induced under oxidative stress conditions, exposure to antibiotics and under host-like conditions^28–30^ and *natT* mutants show reduced survival during antibiotic treatment and in macrophages^30^. While this indicated that RES-domain TA systems are involved in cellular homeostasis and stress response, the mechanisms controlling their activity remain unknown.

Here, we show that an active variant of NatT confers strongly increased tolerance to *P. aeruginosa.* We show that NatT is an NAD phosphorylase, which leads to depletion of both NAD and NADP in a subpopulation of cells. Importantly, while actively growing *P. aeruginosa* cells overcome toxin-mediated NAD deficiencies by inducing the NAD salvage pathway, NatT generates dormant, NAD-depleted cells under nutrient-limited conditions, which spawn hyper-tolerant persisters during outgrowth. Supplementing the growth medium with the NAD precursor nicotinamide blocks toxin expression and activation and eliminates persister formation. Structure-function analyses of the NatR-NatT complex show how NatT interacts with its cognate partner NatR, indicating how this interplay leads to catalytic activation and autoregulation of the NatT toxin. These studies identify a TA system in *P. aeruginosa* that can drive persister formation by modulating a key metabolite of energy metabolism.

## Results

### The NatT toxin induces persister formation in *P. aeruginosa*

We first deleted the *natR and natT* genes on the *P. aeruginosa* chromosome and found that this did not affect growth or fitness under laboratory conditions and did not change survival in the presence of different classes of antibiotics (**Fig. S1a-c**). In contrast, ectopic expression of *natT* from a plasmid increased *P. aeruginosa* survival during drug treatment without affecting growth rates (**Fig. 1b,c**). Tuning *natR* and *natT* expression individually from different inducible promoters, gradually compromised growth and boosted drug tolerance, with increased NatT levels being countered by increased NatR (**Fig. 1d-e**). Ectopic expression of *natT_E29D_*, a mutant isolated in a screen for increased tolerance in *P. aeruginosa*^5^, showed stronger interference with growth (**Figs. 1b**) and increased survival during exposure to tobramycin or ciprofloxacin (**Figs. 1c; S1d**). Thus, the E29D mutation boosts toxin-associated phenotypes, suggesting that it unleashes NatT activity.

To investigate how NatT influences *P. aeruginosa* growth and physiology, we replaced the chromosomal *natT* copy with *natT_E29D_*. The resulting mutant showed normal growth in complex media, but reduced survival in stationary phase, reduced fitness and extended lag periods when stationary cells were diluted into fresh media (**Figs. 1f; S1a**). Diluting stationary phase cultures into fresh media containing tobramycin or ciprofloxacin, showed up to 10’000-fold increased survival compared to wild type (**Fig. 1g**), while resistance levels were unaltered (tobramycin: 1 μg/ml (wt); 1.5 μg/ml (*natT_E29D_*); ciprofloxacin: 0.125 μg/ml (wt); 0.0625 μg/ml (*natT_E29D_*)). Increased expression of *natT_E29D_* gradually lowered growth rates, whereas survival rates reached maximal levels already at expression levels that did not compromise growth (**Fig. S1e-f**). NatT-mediated tolerance was low in rapidly growing cells but strongly increased when cells entered stationary phase (**Fig. 1h**), indicating that NatT-mediated tolerance is most coupled to nutrient limitations.

### NatT is an NAD^+^/NADP^+^ phosphorylase

While expression of NatT in *E. coli* failed to yield soluble protein, combined expression with NatR produced a soluble but inactive NatR-NatT complex. We thus purified the toxin directly from *P. aeruginosa* by affinity chromatography using strains expressing functional FLAG-tagged versions of NatT. While both NatT and NatT_E29D_ were copurified with NatR (**Fig. S2a**), only the complex containing NatT_E29D_ showed activity. Purified NatR-NatT_E29D_ rapidly degraded NAD^+^ generating ADP-ribose-1’’-phosphate (ADPR-1P) and nicotinamide (NAM) as final products (**Figs. 2a-b; S2b**). In accordance with the production of ADPR-1P, NAD^+^ degradation was only observed in presence of phosphate (**Fig. S2b**), demonstrating that NatT is a NAD^+^-dependent phosphorylase. Likewise, purified NatR-NatT_E29D_ was able to degrade NADP^+^ (**Fig. S2c**).

**Figure 2.**
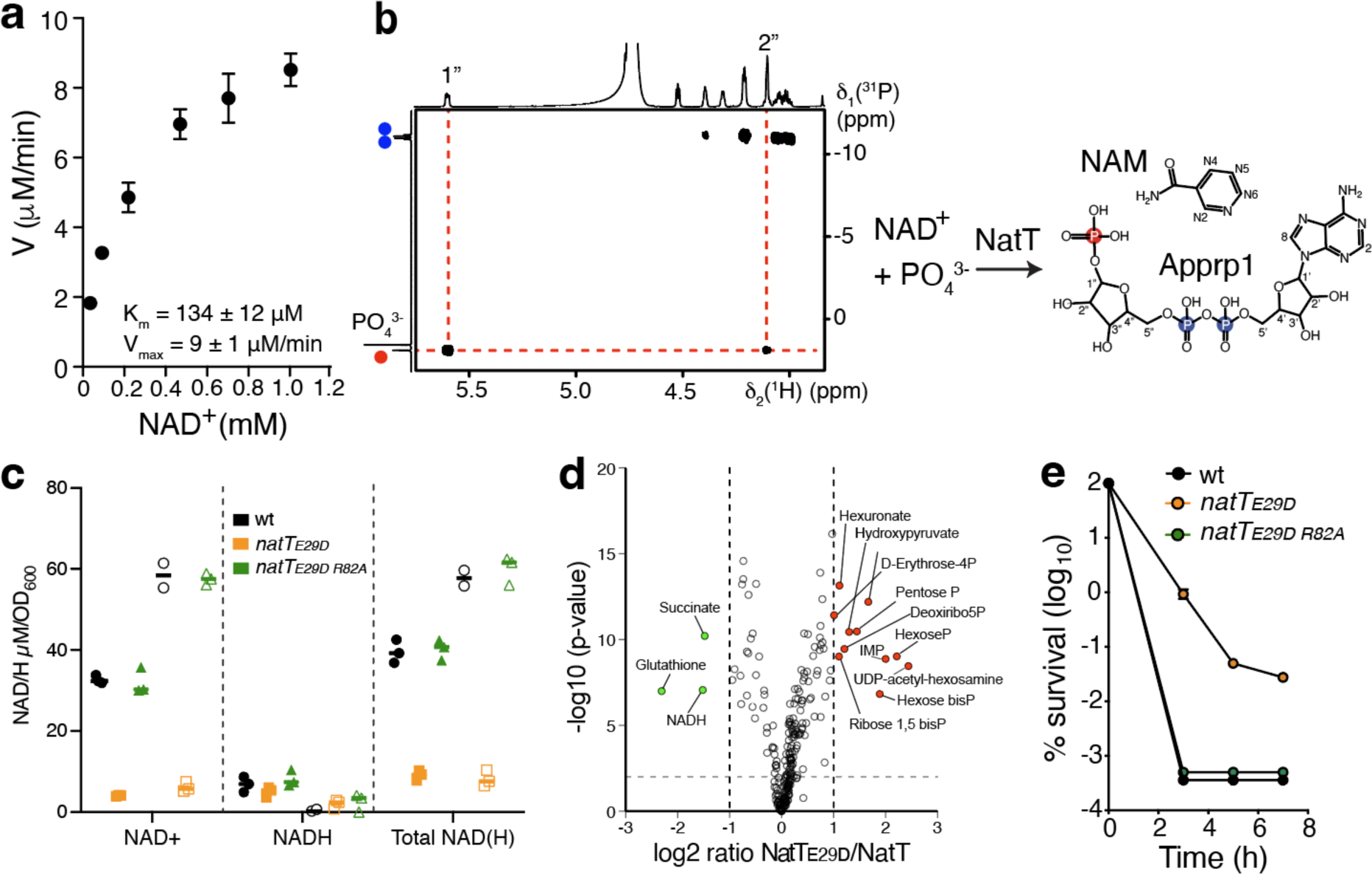
NatT is a NAD-dependent phosphorylase. **a)** Kinetics of NAD^+^ phosphorolysis using purified NatR-NatTE29D complex (50 nM). The kinetic of the reaction was determined by quantifying the NAD^+^ concentration over-time in a series of 1D ^1^H NMR spectra. Km and Vmax values were determined by nonlinear regression analysis with the Michaelis-Menten equation (average ± SEM, n=3). **b)** 2D ^1^H^31^P HMBC NMR NMR spectrum identifying Appr1p as the reaction product of NAD^+^ (5 mM) with 40 nM purified NatRTE29D complex. Phosphate atoms from the ADP-ribose moiety are marked in blue and the phosphate derived from an added orthophosphate group is in red. **c)** NatT depletes cellular NAD^+^ pools of *P. aeruginosa*. NAD^+^ and NADH concentrations were determined in *P. aeruginosa* wild type (wt) and in strains harboring plasmid-encoded copies of *natTE29D* and *natTE29D R82A* growing exponentially (filled bars) or in stationary phase (open bars) (average ± SEM, n≥2). **d)** Expression of *natTE29D* induces global changes of *P. aeruginosa* metabolites. Comparison of metabolites in strains harboring a plasmid with an IPTG-inducible copy of *natTE29D* or a control plasmid. Metabolites significantly enriched or depleted in the *natTE29D* strain are shown in red and green, respectively (n=3). **e)** NatT toxin activity is required for *P. aeruginosa* drug tolerance. Survival of different *P. aeruginosa* strains is shown during exposure to tobramycin (16 μg/ml) (average ± SEM, n>3).

To investigate if NatT depletes NAD^+^ and NADP^+^ *in vivo*, concentrations of NAD^+^/NADH and NADP^+^/NADPH were determined in cultures of *P. aeruginosa* containing plasmid-driven copies of *natT* or *natT_E29D_* harvested in exponential or stationary phase. Both dinucleotides were strongly reduced in cultures expressing *natT_E29D_* compared to wild type (**Figs. 2c; S2e**). Expression of *natT_E29D_* also reduced cellular pools of glutathione and succinate, while intermediates of the pentose-phosphate pathway were increased (**Fig. 2d**). Thus, NatT toxin activity appears to cause redox imbalances, which in turn trigger compensatory metabolic reactions. NAD^+^ and NADP^+^ depletion required NatT catalytic activity, as expression of NatT_E29D R82A_, lacking one of the catalytic core residues of **R**ES domains (see below), did not alter cellular levels of NAD^+^ or NADP^+^ (**Figs. 2c, S2e**), did not affect growth (**Fig. S2f**) and failed to confer drug tolerance (**Fig. 2e**). From this, we concluded that NatT is a NAD^+^/NADP^+^ phosphorylase, which leads to the depletion of both cofactors and to dynamic changes in *P. aeruginosa* metabolism.

### Interaction of NatR and NatT mediates toxin activity

To elucidate the molecular details of NatT activity and its control by NatR, we solved the crystal structure of the NatR-NatT complex. The complex adopts a hexameric structure with two central NatT molecules flanked by two dimers of NatR (NatR’ and NatR”) (**Fig. 3a**). Consistent with the hexameric structure, SEC-MALS analysis revealed a constant mass of 112 kDa in solution (**Fig. S3a**). NatT is composed of a peripheral ring of α-helices and two central β-sheets that form a cavity (**Fig. S3b**). Although NatT is structurally similar to other RES domain proteins^12,14,20^ it harbors an additional structural element corresponding to a 39-residue insertion forming a helix-loop-helix motif (hereafter referred to as ‘Flap’), which provides most of the interaction surface between NatT and NatR’ (**Figs. 3b-c; S3b-d**). The Flap contains several conserved negative charges (E26, D38, E42 and E47), which form electrostatic interactions with positive residues in helices α1, α4, and α7 of NatR’ (**Fig. 3c-d**). For example, E29 of NatT forms an electrostatic interaction with R119, the last but two amino acids of the C-terminal helix α7 of NatR’ (**Fig. 3e**). Intriguingly, the Flap was inserted at the same position of the RES domain fold multiple times independently during evolution (**Fig. S3d-e**), indicating that it has adopted a role in RES toxin control by modulating toxin-antitoxin interaction. In line with this idea, the activating mutation E29D is located precisely at the hinge between the NatT Flap and helix α7 of NatR’, which controls access to the catalytic cavity of NatT (**Figs. 3e; S3d**). Moreover, a well-defined tetrahedral atomic density modeled as a phosphate molecule was found in the interface between the Flap and NatR’, in close vicinity of residue E29 (**Figs. 3c,e; S3b**). Residues coordinating the phosphate moiety (R70 and R119 of NatR’; K54, Y66, E104 and Y74 of NatT) are highly conserved in NatR and NatT homologs and form a pocket at the NatT-NatR’ interface (**Figs. 3e; S3d**).

**Figure 3.**
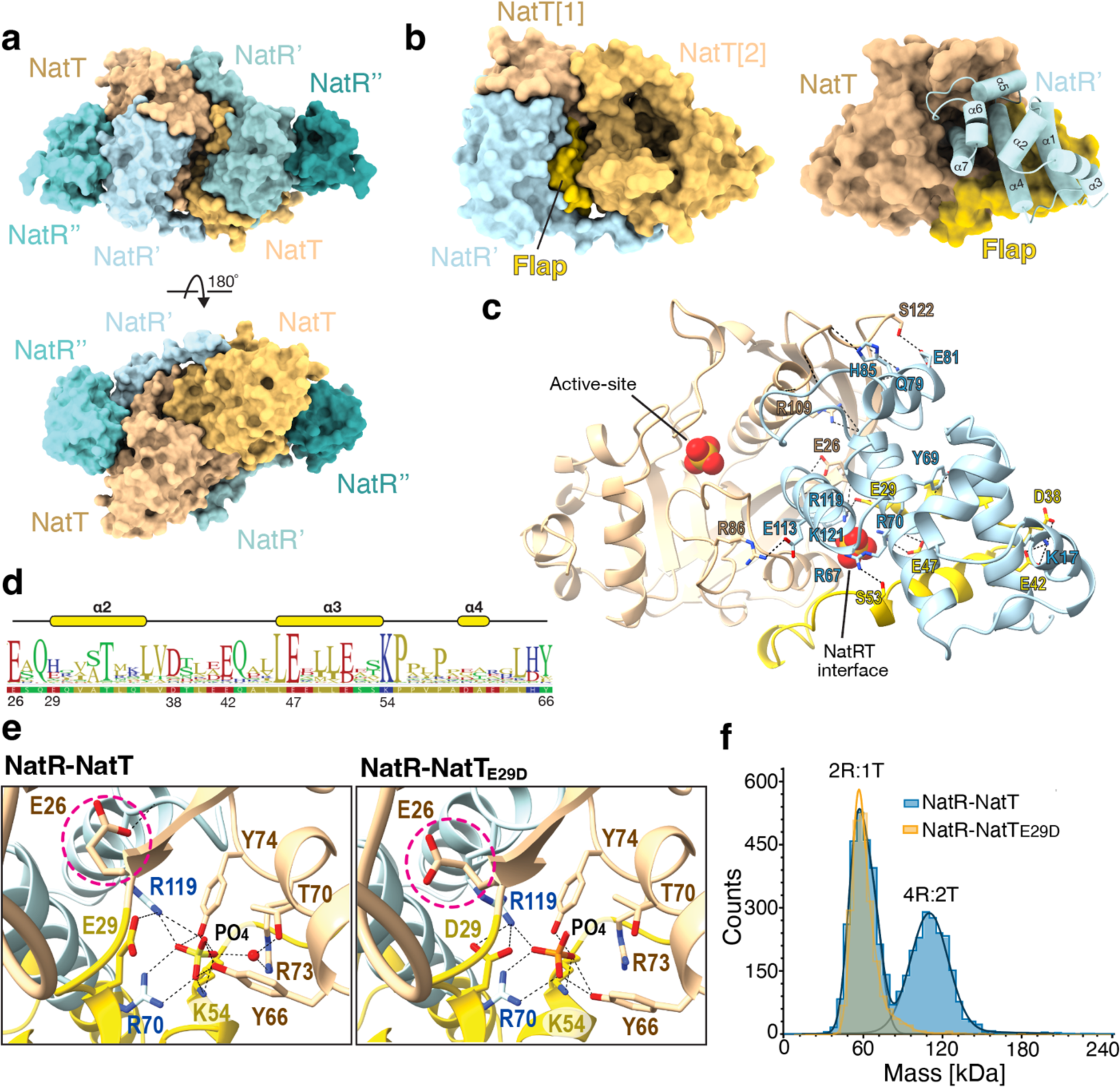
Interaction of NatR and NatT mediates toxin activation. **a)** Crystal structure of a NatR-NatT hexamer with a central NatT dimer and two flanking dimers of NatR (NatR’ and NatR’’). **b)** NatR-NatT interaction is mediated by the Flap region. Left: Surface representation of NatT [protomer 1] with its Flap region (yellow) sandwiched between the interacting NatR’ and the neighboring NatT [protomer 2]. Right: Surface representation of NatT with NatR in cartoon style and individual helices marked. NatR’ interacts with the Flap region of NatT and α7 of NatR’ extends into the active site groove of NatT. **c)** Detailed view of NatT-NatR’ interface with negatively charged residues of the Flap (yellow) and positively charged residues of NatR’ indicated in stick representation. Phosphate moieties in the active site and in the NatR-NatT interface are indicated as space filled molecules. **d)** Conservation of Flap region (NatT residue 26 to 66). Positions of conserved residues interacting with NatR’ are highlighted. K54 and Y66 coordinate the phosphate molecule in the NatR-NatT interface (see: **e**). **e)** Zoom-in views comparing the interaction of NatR’ with NatT wild type and NatTE29D. Residues with altered conformation are highlighted in stick representation with coloring scheme as in (**c**). The movement of residue E26 away from its interaction with the backbone of α7 of NatR’ (blue) is marked by a red circle. **f)** Mass photometry analysis of NatR-NatT (blue) and NatR-NatTE29D (yellow) at 10 nM. The mass of the NatR-NatTE29D complex corresponds to a NatR’-NatR’’-NatT complex.

Superimposing the structures of NatT and diphtheria toxin bound to its NAD^+^ substrate indicated that the active site of NatT is located at the base of the central cavity formed by its β-sheet core^31^. Active site residues are conserved in NatT and other RES domain proteins, including R23, R82, Y93, N193, and S184 (**Fig. S3b,d**)^12^. Residues R82, Y108, Y162, and R186 coordinate a second phosphate molecule in the active site of NatT, positioned next to the nicotinamide group of NAD^+^, that likely participates directly in the cleavage of NAD^+^ (**Fig. S3b**). In the NatR-NatT complex, the C-terminal helix α7 of NatR’ extends all the way into the substrate-binding cavity of NatT (**Fig. 3b-c**), indicating that NatR’ controls toxin activity by blocking substrate access and that dynamic repositioning of α7 and NatR’ is likely required to activate NatT.

To explore the NatT activation mechanism, we next solved the structure of NatT_E29D_ in complex with NatR. Structural differences between wild-type and mutant proteins were limited to subtle changes in the NatR’-NatT interface close to residue E29. A water molecule bridging the phosphate positioned in the NatR’-NatT interface with residues T70 and R73 of NatT wildtype was missing in the E29D mutant (**Fig. 3e**). Moreover, the E29D substitution changes its interaction with R119 of NatR’, forcing the neighboring E26 to adopt a different rotameric configuration, which in turn disrupts the interaction with the main chain of the C-terminal helix α7 of NatR’. Based on this, we speculate that the E29D mutation weakens the interaction between the NatT Flap and the NatR’, thereby promoting the disassembly of the stable hexameric structure. This is in line with SEC-MALS experiments showing that the NatT-NatR wild-type complex is a stable hexamer (112 kDa) at micromolar concentrations, whereas the complex containing NatT_E29D_ displayed an additional peak with significantly lower mass (**Fig. S3a**). Mass photometry analysis at low nanomolar protein concentrations revealed two different oligomeric states for the NatR-NatT wild-type complex, a 112 kDa hexamer and a 60 kDa species, which likely corresponds to a complex between a NatR dimer and NatT (2R:1T) (**Fig. 3f**). Importantly, NatT_E29D_ showed a single peak at 60 kDa, indicating that the NatT-NatR complex is in a dynamic equilibrium between an inert hexameric state and a smaller, active subcomplex and that this transition mediates NatT toxin activation. Of note, the Flap is tightly sandwiched between NatR’ and the adjacent NatT protomer **(Fig. 3b)**, restricting Flap movement and keeping the active site pocket occluded by the C-terminus of NatR’. Thus, disruption of the NatT-NatT dimer could liberate the Flap, allowing it to flexibly move while still bound to NatR’, thereby providing access to the active site.

### NatT activity is coupled to *natR-natT* transcription

When monitoring NatR and NatT protein levels in different *P. aeruginosa* strains, we noticed that both proteins are elevated in the *natT_E29D_* mutant compared to wild type (**Fig. 4a-b**). To examine if this results from altered transcription, we engineered *P. aeruginosa* strains carrying a *gfp* reporter downstream of *natR-natT* on the chromosome (**Figs. 4a; S4a**). Expression of *natT* was strongly upregulated in a Δ*natR* mutant, but reduced to wild-type levels when *natR* was complemented from a plasmid (**Figs. 4c, S4b**). Likewise, in a Δ*natT* mutant transcription of *natR-natT* was derepressed (**Fig. 4c**) and NatR levels strongly upregulated (**Fig. 4d**). Thus, *natR-natT* transcription is autoregulated with both NatR and NatT contributing to its repression. Intriguingly, while *natR-natT* transcription remained at low levels in *P. aeruginosa* wild type, it strongly increased in the *natT_E29D_* background as cells progressed toward stationary phase (**Figs. 4e; S4c**). NatT and NatR protein levels were also elevated in the *natT_E29D_* background, with differences being most prominent in stationary phase (**Figs. 4f; Fig. S4d-e**).

**Figure 4.**
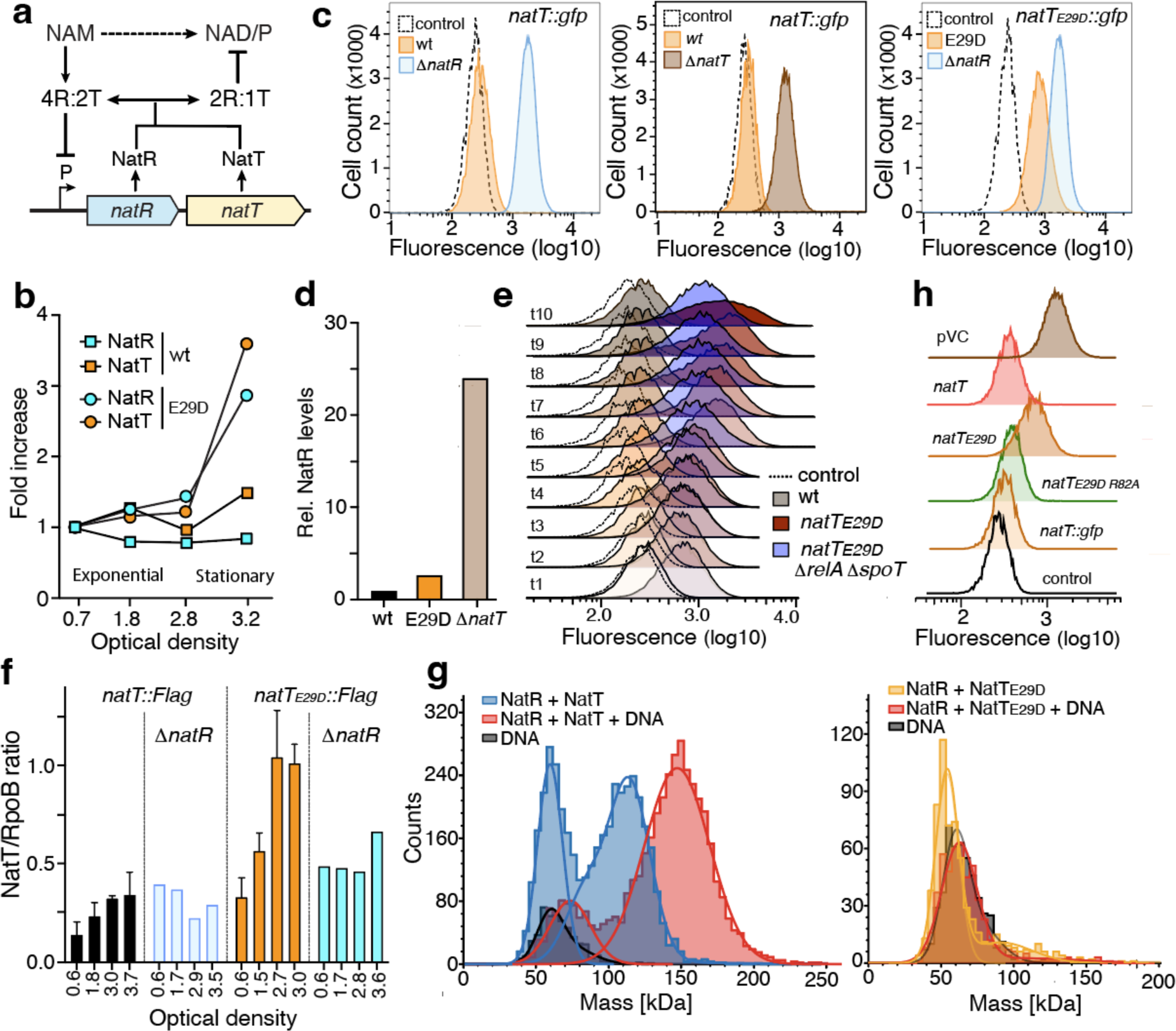
NatR is an antiand a co-toxin of NatT. **a)** Schematic model of NatT activation and autoregulation. A catalytically inactive hexameric complex (4R:2T) acts as repressor of *natR-natT* transcription. Transition to a 2R:1T complex leads to derepression of *natR*-*natT* transcription and under specific conditions to NatT NADase activation. The postulated regulatory role of the NAD precursor nicotinamide (NAM) is indicated. **b)** NatR (turquoise) and NatT (orange) levels are increased in a *natTE29D* mutant. Proteins were quantified by mass spectrometry and plotted as a function of growth (optical density) of *P. aeruginosa* wild type (wt) and *natTE29D* mutant. **c)** Transcription of *natT* is controlled by the NatR-NatT toxin complex. Transcription was determined in reporter strains containing a chromosomal copy of *gfp* downstream of *natT*. Fluorescence of a control strain lacking the *gfp* reporter is indicated (dotted lines). **d)** NatR protein levels are increased in *natTE29D* and Δ*natT* mutants. Levels of NatR were determined by quantitative mass spectrometry analysis of *P. aeruginosa* wild type and mutants indicated. **e)** Transcription of *natR-natT* is induced in the *natTE29D* background. Cultures of *P. aeruginosa* wild type, *natTE29D* and *natTE29D* Δ*relA* Δ*spoT* mutants harboring a chromosomal *natT*::*gfp* reporter were assayed by FACS during different phases of growth (see: Fig. S4c). Black lines show baseline auto-fluorescence of a strain lacking a *gfp* reporter gene. **f)** NatT protein levels are reduced in a strain lacking NatR. Immunoblots of *P. aeruginosa* wild-type and Δ*natR* mutants with chromosomal *natT*-FLAG or *natTE29D*-FLAG alleles. Cell extracts were harvested at different ODs and stained with anti-FLAG and anti-RpoB antibodies and NatT levels normalized to RpoB. **g)** NatR-NatT binds to its own promoter. Mass photometry of NatR-NatT (blue) and NatR-NatTE29D (yellow) without and with DNA (red) containing the *natR* promoter region. The mass of dsDNA is shown in black. **h)** Transcription of *natR-natT* activity is coupled to NatT activity. Expression of a chromosomal *natR-*Δ*natT::gfp* reporter was assayed by FACS in a *P. aeruginosa* Δ*natT* mutant expressing different *natT* alleles from a plasmid, as indicated. The black line represents the fluorescent signal in a control strain lacking a *gfp* reporter. One representative experiment is shown (pVC=plasmid control).

The above findings indicated that transcription of *natR*-*natT* is coupled to the activity of the NatT toxin. In agreement with this, purified NatR-NatT wild-type complex (4R:2T) readily bound to the *natR*-*natT* promoter region, while a purified NatT_E29D_ complex failed to bind the same DNA fragment (**Fig. 4g**). Moreover, while *natR-natT* transcription was derepressed in the *natT_E29D_* background, it remained fully repressed in a *natT_E29D R82A_* mutant, encoding a catalytically inactive NatT toxin (**Fig. 4h**). Together with the observation that NatT_E29D_ failed to form a stable hexameric complex with NatR *in vitro* (**Fig. 3f**), these findings indicated that NatT toxin activity and *natR-natT* expression are controlled through a dynamic equilibrium between a stable NatR-NatT hexamer and a smaller NatR-NatT complex that is unable to adopt the conformation needed to bind to the *natR*-*natT* promoter region (**Fig. 4a**). This model is supported by the observation that increasing cellular levels of NatT not only increased autotoxicity and drug tolerance (**Fig. 1d,e**) but also led to the progressive activation of the *natR-natT* promoter (**Fig. S4f**), presumably by shifting the NatR-NatT equilibrium and by that modulating the hexameric repressor conformation.

Despite of *natT* transcription being derepressed in strains lacking NatR, NatT activity was completely abolished in a Δ*natR* mutant (**Fig. 1d,e**). This indicated that NatR acts as anti-toxin blocking NatT expression and activity, and at the same time, is required for toxin function. In line with this, NatT protein levels remained low in a Δ*natR* mutant (**Fig. 4f**), most likely due to its poor solubility and rapid degradation in the absence of NatR (**Figs. S4g-h**). Deleting *natR* abolished NatT toxin activity (**Fig. S4i**) as well as NatT-mediated drug tolerance (**Fig. S4j-l**). From this, we concluded that NatR serves as anti- and co-toxin for NatT and that toxin regulation is mediated by subtle changes in NatR-NatT interaction and complex stoichiometry, a process that strictly requires NatR.

### NatT-mediated antibiotic tolerance is countered by the NAD salvage pathway

*P. aeruginosa* replenishes its NAD^+^ pool via *de novo* synthesis from aspartate or, if available, from nicotinamide (NAM) through the NAD^+^ salvage pathway I (NSPI) (**Fig. 5a**)^32^. Importantly, the NAD salvage pathway was recently shown to be essential for *P. aeruginosa* growth under conditions mimicking human infections^33^. Deletion of NSPI genes *pncB*, *nadD2* and *nadE* strongly increased NatT-mediated toxicity (**Fig. 5a**), indicating that the NAD^+^ salvage pathway balances NAD^+^ shortages in cells inducing NatT toxin activity. To test this, we engineered a strain carrying a *gfp* reporter downstream of *pncB1* on the *P. aeruginosa* chromosome and showed that deletion of *nrtR*, the gene encoding the NSPI repressor^32^, led to strong derepression of *pncB1* (**Fig. 5a-b**). Moreover, *pncB1* was derepressed in a small subpopulation of the n*atT_E29D_* mutant but not of *P. aeruginosa* wild type (**Fig. 5b-c**). Cells inducing NSPI genes were not observed in a *natT_E29D R82A_* background, demonstrating that derepression of the NAD^+^ salvage pathway indeed requires an active NatT toxin (**Fig. 5c**). Increased expression of *natT_E29D_* increased the fraction of cells with derepressed NSPI genes, while increasing *natR* expression had the opposite effect (**Fig. S5a-c**). Time-lapse microscopy of the *natT_E29D_* mutant with the *pncB1-gfp* reporter revealed that cells stochastically inducing the NAD^+^ salvage pathway transiently slowed growth but quickly resumed concomitant with the loss of the fluorescent signal. Thus, under nutrient-rich conditions, derepression of the NAD^+^ salvage pathway effectively neutralizes stochastic NatT activation. Intriguingly, the fraction of cells inducing NSPI genes gradually declined during late log phase and upon entry into stationary phase, indicating that the metabolic response of *P. aeruginosa* to NatT activity collapses as nutrients become limited (**Fig. 5c**).

**Figure 5.**
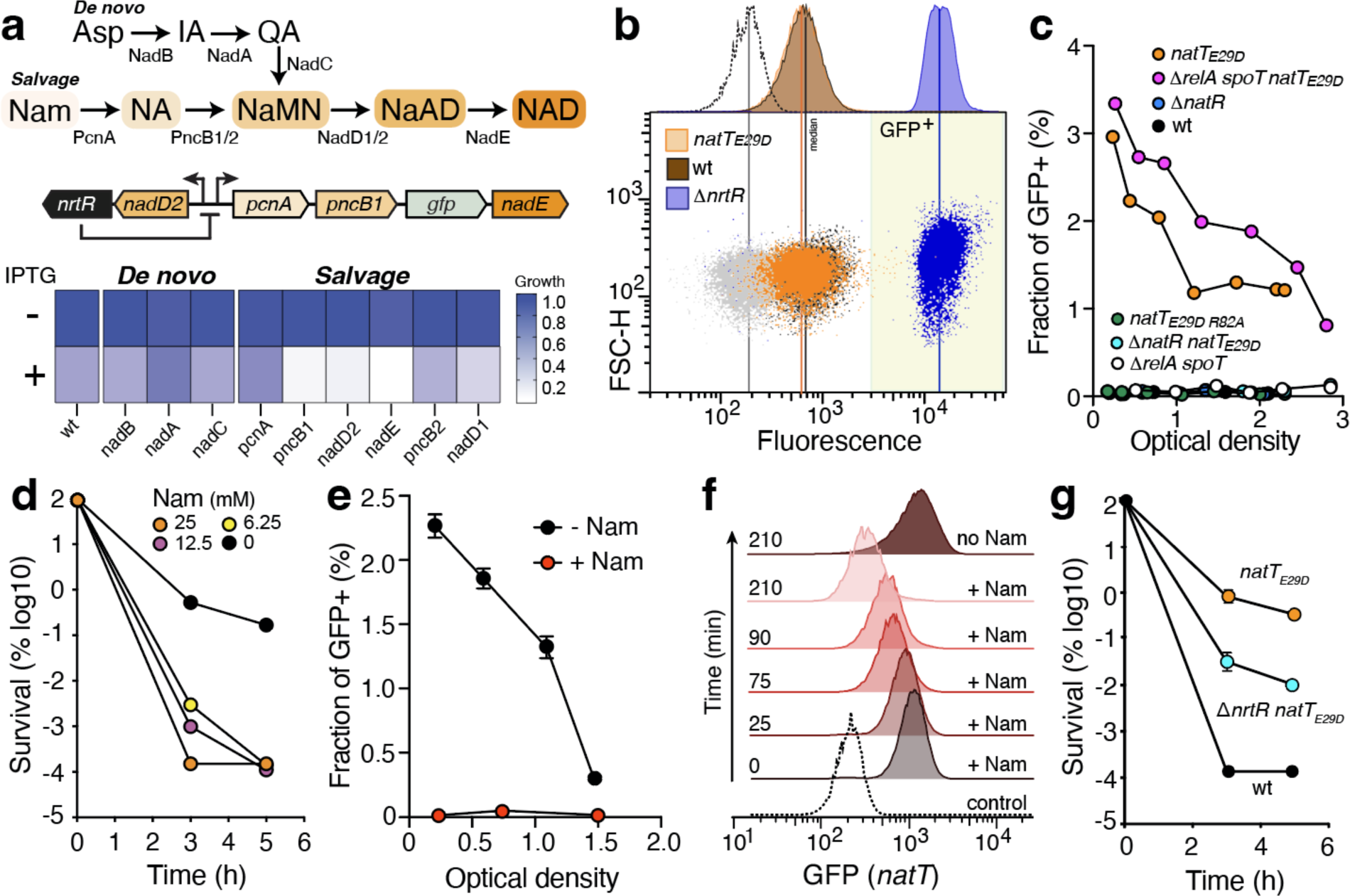
The NAD salvage pathway neutralizes NatT toxin activity and abolishes drug tolerance. **a)** Schematic of NAD^+^ *de novo* and salvage pathways in *P. aeruginosa* (top) with intermediates and catalyzing enzymes indicated. Salvage pathway regulation by NrtR is shown below with enzymes and genes being highlighted in matching colors. The position of the chromosomal *gfp* reporter used for these studies is in green. Impact of *natTE29D* expression on growth of *P. aeruginosa* wild type and different salvage pathway mutants is shown on the bottom. Strains carrying an IPTG-inducible copy of *natTE29D* on a plasmid were grown in LB medium with or without IPTG. The heat map shows the OD after 8 hours of growth normalized to the OD of wild type. **b)** Transcription of NAD salvage pathway genes in individual cells of *P. aeruginosa* wild type, *natTE29D*, and Δ*nrtR* mutants harboring a *pncB1*::*gfp* reporter was analyzed by FACS. GFP signals are shown for individual cells and as histograms. A strain lacking the *gfp* reporter is shown as control (grey dots and stippled line). The area with average GFP signals higher than wild type (GFP^+^) is marked in yellow. **c)** NatT-mediated activation of salvage pathway genes declines in a growth phase-dependent manner. Fractions of cells expressing salvage pathway genes are shown as a function of the optical densities of cultures analyzed. **d)** Nicotinamide (NAM) abolishes NatT-mediated drug tolerance. *P. aeruginosa natTE29D* grown in LB supplemented with different concentrations of NAM was challenged with tobramycin (32 μg/ml) for 3 hrs and fractions of surviving cells were determined. **e)** NAM abolishes NatT-mediated NAD^+^ salvage pathway induction. Fractions of cells with induced NAD^+^ salvage pathway were determined as in (**c**) in the presence or absence of NAM (20 mM) (average ± SEM, n=2). **f)** NAM blocks NatT-mediated derepression of the *natR-natT* operon. Growing cultures of the *natRTE29D*::*gfp* reporter strain were supplemented with NAM (25 mM) and analyzed by at times as indicated. A control strain lacking a *gfp* reporter is shown (dotted line). **g**) Salvage pathway derepression partially abolishes NatT-mediated drug tolerance. Experiments were carried out as indicated in (d).

To assess the impact of NatT on *P. aeruginosa* growth, we engineered reporter strains constitutively expressing the fluorescent protein TIMER^34^, which directly reads out growth rates of individual cells without compromising NatT-mediated persister formation (**Fig. S6a**). While growth was not compromised in wild type and the *natT_E29D_* mutant during the logarithmic growth phase, numbers of growth-arrested cells spiked up in the mutant and, to a lower extent in wild type, upon nutrient depletion (**Figs. 6a, S6b**). In this experiment, samples from stationary phase were diluted into fresh medium and TIMER-based fluorescence was analyzed after three hours of growth resumption. Spotting stationary cultures directly onto agar patches containing fresh nutrients showed that wild-type cells resumed growth immediately, while a large fraction of *natT_E29D_* mutant cells indeed showed lag phases of variable lengths (**Fig. 6b**). Thus, NatT activity in stationary phase generates arrested cells that experience extended lag periods when exposed to fresh nutrients.

**Figure 6.**
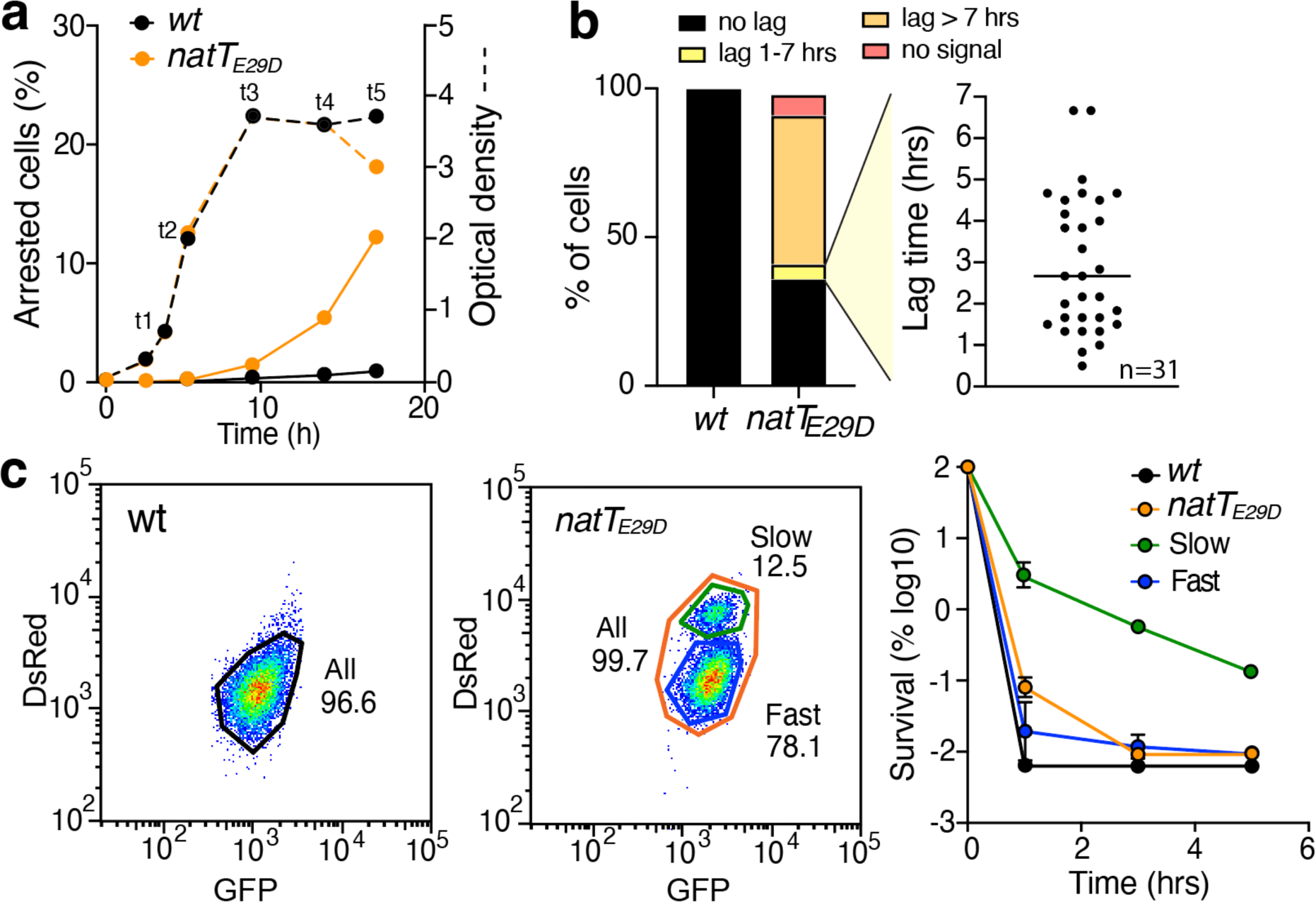
NatT activation generates persister subpopulations with extended lag phase. **a,b)** The *P. aeruginosa natTE29D* mutant generates growth arrested cells. Cultures of *P. aeruginosa* wild type and *natTE29D* mutant expressing TIMER were grown in LB and samples removed at indicated the time points (t1-t5), diluted into fresh medium (OD 0.1) and analyzed by FACS after three hours of growth. Fractions of arrested cells are plotted in (**a**) (solid lines). **b)** Cells harvested at time point t5 (**a**) were spotted on LB agar patches and their growth analyzed by time-lapse microscopy (wild-type, n=450; *natTE29D*, n=150). Bar plots indicate fractions of cells with delayed re-growth and different lag times (black line=median). **c)** The *natTE29D* mutant generates growth-arrested cells with increased drug tolerance. Stationary phase cultures of wild type and *natTE29D* mutant expressing TIMER were harvested at time point t5 **(a)**, diluted into fresh LB medium for three hours, and sorted by FACS using the red and green fluorescence channels (left panels). Subpopulations were harvested and survival was determined upon exposure to tobramycin (right panel).

To investigate if the observed transient growth arrest contributes to antibiotic survival, stationary phase cultures of strains expressing TIMER (**Fig. 6a**) were resuspended into fresh media for 3 hours, followed by FACS sorting to separate growing from arrested subpopulations using green and red fluorescence (**Fig. 6c**). Exposing sorted populations to high concentrations of tobramycin (16 μg/ml) revealed rapid killing of *P. aeruginosa* wild type and the *natT_E29D_* subpopulation that showed rapid growth recovery. In contrast, the subpopulation with slow growth recovery showed significantly increased survival (**Fig. 6c**). From this, we concluded that stochastic activity of NatT leads to NAD^+^ depletion in a subpopulation of cells, which in turn provides drug tolerance during the outgrowth from stationary phase.

To further scrutinize this model, we supplemented the growth medium with NAM, the precursor of the NAD^+^ salvage pathway (**Fig. 5a**). NAM not only reduced NatT-mediated drug tolerance in a concentration-dependent manner (**Fig. 5d**), but also abolished NatT-mediated derepression of the NAD^+^ salvage pathway (**Fig. 5e**), restored growth of strains ectopically expressing *natT_E29D_* from a plasmid (**Fig. S5d**), alleviated NatT-mediated hyper-sensitivity of an NAD^+^ salvage pathway mutant (**Fig. S5e**), and reduced the fraction of growth arrested cells during outgrowth from stationary phase (**Fig. S5f**). NAM supplementation also effectively blocked *natR*-*natT* transcription in the *natT_E29D_* mutant (**Fig. 5f**) and reduced levels of NatR and NatT proteins in the same strain (**Fig. S5g**). Finally, genetic derepression of salvage pathway genes in an λι*nrtR* mutant (**Fig. 5a**) greatly reduced NatT-mediated persister formation (**Fig. 5g**).

The observation that NAM neutralized the physiological consequences of NatT even in the absence of a functional salvage pathway (**Fig. S5e**), indicated that it abolished persister formation by interfering with toxin expression and/or activity rather than by simply replenishing pools of nicotinamide dinucleotides. A possible candidate mediating NAM-specific effects is the global alarmone (p)ppGpp, which controls bacterial physiology under nutrient-deplete conditions^35^ and was implicated in antibiotic tolerance^36–38^. To test this, we introduced the *natT_E29D_* allele into a strain lacking RelA and SpoT, the enzymes responsible for (p)ppGpp synthesis^39^. While NatT-mediated tolerance was indeed completely abolished in a strain lacking (p)ppGpp (**Fig. S6c-d**), this effect was not due to interference with NatT activity as both *natR*-*natT* transcription (**Fig. 4e**) and derepression of the NAD^+^ salvage pathway (**Fig. 5c**), were unchanged in the λι*relA* λι*spoT* mutant. Thus, (p)ppGpp does not promote *P. aeruginosa* drug tolerance by regulating NatT toxin activity, but rather by controlling downstream processes that help *P. aeruginosa* cope with the physiological consequences of NatT-mediated NAD^+^ depletion.

### NatT is under positive selection during infections in humans

In order to assess the potential role of NatR-NatT during *P. aeruginosa* infections, we analyzed *natR* and *natT* gene sequences in 8286 genomes of *P. aeruginosa* available in the SRA database^40^. While NatR sequences are highly conserved, more than half of the strains analyzed contained single or multiple SNPs in NatT (**Fig. 7a**). The most frequent changes were I14V and V117A, which occurred as single SNPs or in combinations with other mutations and likely mark the PA14 strain lineage. Additional *natT* SNPs were found in 67% of all isolates from CF patients (n=479), 83% of the isolates from diverse acute infections (n=161), and 80% of other hospital isolates (n=305) (**Table S1**), arguing that NatT toxin function is under selective pressure during human infections. The E29D mutation was not found in any of these strains, indicating that high fitness costs may prevent its selection *in vivo* (**Fig. S1a**). A selection of the most frequent *natT* alleles was amplified from patient isolates and grafted into *P. aeruginosa* lab strain PAO1. While none of these alleles showed a toxic effect when expressed ectopically, survival rates gradually increased with increasing numbers of *natT* mutations, with one variant harboring five replacements conferring the highest level of tolerance (**Fig. 7b**) and activation of the NSPI salvage pathway genes similar to the *natT_E29D_* allele (**Fig. 7c**). Of note, this *natT* allele carried mutations in residues S105 and P165, which are positioned at the NatR-NatT interface, close to the active site of NatT.

**Figure 7.**
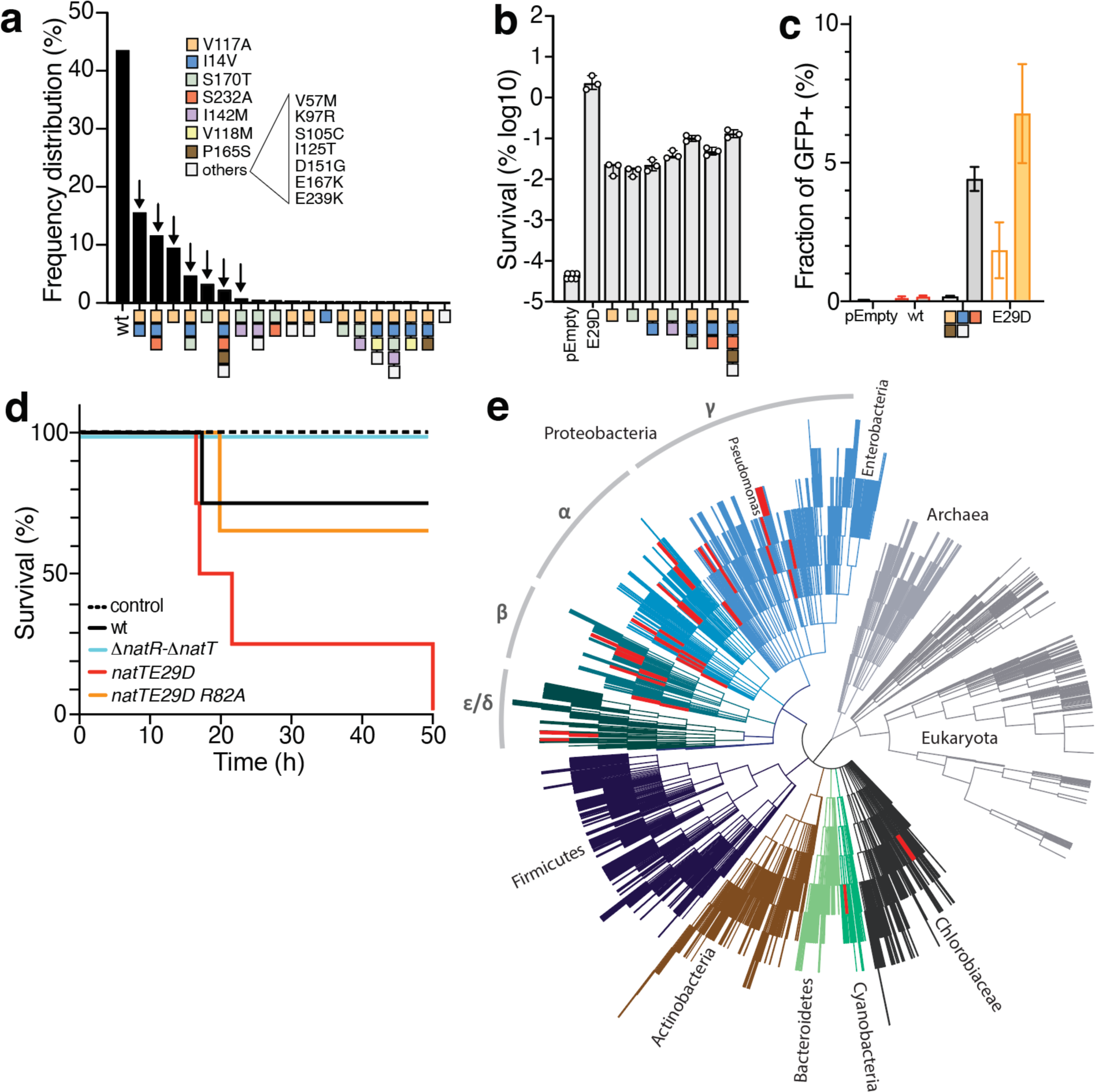
The NatRT module is widespread in proteobacteria and is under selection during infections. Frequency distribution of *natT* alleles in 8286 *P. aeruginosa* strains analyzed. Single SNPs and SNP combinations are indicated in different colors. Arrows indicate the *natT* alleles that were tested in this study. **(b)** Different *natT* alleles confer different levels of drug tolerance. Cultures of *P. aeruginosa* carrying plasmids with different *natT* alleles from **(a)** were scored for survival after 3 hrs of treatment with tobramycin. **(c)** Salvage pathway induction by different *natT* alleles. A *P. aeruginosa pncB1*::*gfp* reporter strain carrying plasmids with different *natT* alleles under IPTG control was grown with (filled bars) or without (open bars) IPTG. Fractions of cells with derepressed salvage pathway (GFP^+^) are indicated. **d)** NatT activity controls *P. aeruginosa* virulence in a simple insect larvae model. Survival rates of *G. mellonella* larvae are shown for different *P. aeruginosa* strains as indicated. Experiments were carried out with five larvae per strain tested. **e)** Bacterial phylogeny with the distribution of NatR-NatT TA system orthologs (red bars).

These observations suggested that mutations activating NatT accumulate during infections in humans, possibly increasing stress tolerance under these conditions while balancing fitness costs. Alternatively, NatT-mediated NAD depletion and salvage pathway induction could generate a *P. aeruginosa* subpopulation with altered virulence. NrtR, the repressor of NSPI salvage pathway genes, also regulates virulence gene expression, including T3SS^41^ and T6SS^42^. To test this possibility, we used different *natT* mutants to infect *Galleria* larvae, a simple non-vertebrate infection model for *P. aeruginosa*^43^. Intriguingly, a *natT_E29D_* mutant showed strongly increased killing, while survival of larvae infected with a 11*natT* mutant was increased (**Fig. 7d**). These findings are compatible with the idea that NatT toxin activation stimulates both *P. aeruginosa* virulence and persistence.

## Discussion

Self-intoxicating degradation of cellular pools of NAD/NADP is used by bacteria to defend against bacteriophages in a process called abortive infection^15–18^. But while known bacterial immunity systems targeting NAD exclusively adopt TIR and SIR2 NADases^19^, the physiological role of RES domain toxins has remained unclear^12,13^. Here, we show that an activated variant of NatT, a RES domain toxin that is part of the core genome of *P. aeruginosa*, strongly promotes survival during treatment with different classes of bactericidal antibiotics. NatT belongs to a subgroup of RES domain proteins that have adopted a Flap region mediating its interaction with the antitoxin NatR. Orthologs of the NatR-NatT TA system are primarily found in proteobacteria, where they seem to have spread horizontally to distinct clades in alpha-, beta-, gamma-, deltaand epsilon-purple bacteria, but are absent in enterobacteria (**Fig. 7e**). This wide taxonomic distribution and the observation that none of the *natR* or *natT* homologs are associated with phage genes or phage remnants, argues against NatT playing a role in phage defense. In line with this, probing a selection of more than 50 natural *P. aeruginosa* phage isolates has failed to show differential susceptibility of wild type and a 11*natT* mutant.

Although TA systems have been implicated in persister formation in response to stress, the idea that they contribute to fitness of bacterial populations under adverse conditions has remained controversial^1,44–46^. Stress-mediated phenotypes of TA null mutants are often missing^47–50^ and transcriptional upregulation of TA systems in response to diverse stressors did not result in active RNAse toxins in *E. coli*^51^. In agreement with this, we found that even in strains that globally depressed *natR-natT* transcription, NatT activity remained limited to a small subpopulation, arguing that additional, post-transcriptional levels of NatT control must exist and that additional signals may be required for toxin activation. Limiting toxin activity to a small subpopulation may also explain why stress-mediated expression of TA systems in *E. coli* does not result in observable growth inhibition^51^. Moreover, our data showed that even when NatT is unleashed, actively growing cells can neutralize its activity by replenishing NAD. Whereas NatT-dependent derepression of NAD salvage pathway genes effectively evaded growth obstruction, salvage pathway mutants were hypersensitive to NatT. The observation that the fraction of cells with derepressed salvage pathway gradually declines during entry into stationary phase, indicated that NatT-mediated NAD/P depletion can no longer be countered when cells run out of nutrients, thereby establishing a population of low NAD persister cells. This explains why NatT-mediated persister formation requires stationary phase physiology, while actively growing cells show little drug tolerance. Thus, NatT activity together with differential regulation of salvage pathway components could adjust the formation of persisters when bacteria experience nutritional stress.

*P. aeruginosa* harbors a large variety of type II TA systems located on the chromosome, plasmids or prophages^52^. However, only four of these, including NatT, are part of the core genome^53^. Similar to other RES domain toxins^12^, *natT* expression is induced by different stressors, including redox stress, antibiotics or in media mimicking host conditions^28–30^. Moreover, a *P. aeruginosa natT* mutant was recently shown to be more susceptible to killing by different classes of bactericidal antibiotics and to promote survival in macrophages^30^. This is reminiscent of the proposed role for the RES domain toxin MbcT in survival of *M. tuberculosis* in macrophages^12^. A specific role in host persistence could also explain our findings that NatT modulates host killing in a simple insect larvae infection model. Thus, although the physiological cues regulating NatT expression and activity remain to be defined, this TA system seems to play an important role in *P. aeruginosa* homeostasis and fitness in the host. In line with this, we have identified *natT* alleles in patient isolates displaying mild toxin activation and increased antibiotic tolerance. It is possible that NatT function is under selection during infections to increase basal level persistence provided by this TA system.

NatT function is tightly controlled by NatR, which is both an antitoxin and an auxiliary factor required for toxin stability. Genetic, biochemical and biophysical analyses suggested that the NatR-NatT complex is able to switch between an inert hexameric state 4R-2T and an active 2R-T complex. Because only the hexameric complex is able to bind DNA, we propose that NatT activity is coupled to transcriptional autoregulation through this oligomeric switch. The NatT Flap provides a large interaction surface for the antitoxin, effectively locking NatR’ and its C-terminal helix in the NatT active site cavity. NatT activation likely includes the disintegration of the NatT dimer, which in turn could liberate the Flap and allow NatR’ to move away from the active site cavity without disengaging from its toxin partner. Residue E29 interacts with the C-terminal helix α7 of NatR’ and, through its immediate neighbor Q30, also participates in NatT dimerization. Thus, the activating mutation E29D may not only disturb the interaction between the Flap and helix α7, but may also disrupt NatT dimerization. If and how the phosphate molecule positioned at the Flap-NatR’ interface influences NatR-NatT interaction, remains to be shown. Residues coordinating this phosphate are highly conserved, arguing that it has likely adopted an important structural or mechanistic role. Although this pocket is not accessible in the fully closed hexameric state, it is possible that rearrangement of Flap and NatR’ in the active conformation provides access for phosphorylated metabolites that regulate NatT activity. A possible candidate is ADPR-1P, one of the NAD breakdown products generated by NatT. Positive feedback mediated by this unique metabolite could explain why a catalytic NatT mutant (R82A) was unable stably derepress *natR-natT* transcription. In agreement with NatT and NatR forming a stable interaction through the Flap, NatT is rapidly degraded in *P. aeruginosa* strains lacking NatR and, in contrast to RES domain toxins lacking a Flap^12,20,54^, showed no toxicity by itself or when expressed in *E. coli*. The Flap region was adopted by RES domain toxins several times independently (**Fig. S3e,f**), arguing that it marks an important evolutionary step towards more sophisticated toxin control by their cognate antitoxins.

Although this work has revealed exciting new mechanistic insights, the signals or physiological cues responsible for NatT toxin activation and its downstream consequences remain to be defined. Several important aspects need attention. 1. NatT toxin activity was limited to a small subpopulation of cells even in strains constitutively expressing *natR* and *natT*. Which auxiliary factors or mechanisms are responsible for this stochastic control? 2. Derepression of *natR*-*natT* transcription requires NatT catalytic activity. How is NAD degradation coupled to *nat* gene expression? 3. Supplementing media with the NAD precursor nicotinamide effectively blocked NatT activity and expression. How do metabolic cues feedback on the NatR-NatT system and could this help explain its stochastic nature? While this effect could be mediated indirectly, it is also possible that NatT is directly inhibited by its product, similar to the mammalian TIR domain NADase SARM1, a central switch in axon degeneration^55^. 4. Similar to MbcT from *M. tuberculosis*^12^, NatT is an NAD phosphorylase converting NAD into Nam and ADPR-1P. ADPR-1P is known to be produced during tRNA splicing^56^ in eukaryotes, but has not been implicated in any other cellular processes. Does ADPR-1P have a specific role in mediating NatT downstream effects distinct from ADP-ribose (ADPR), the breakdown product of physiological NAD hydrolases? And does the salvage pathway repressor NrtR distinguish between ADPR-1P and its cognate ligand ADPR^32^? TIR domain NADases involved in cell death signaling and abortive infection produce cyclic derivatives of ADP-ribose as potent signaling molecules^18,57^. It will be interesting to test if ADPR-1P has similar properties or if RES domain phosphorylases operate on the simple logic of depleting NAD^+^ pools.

## Acknowledgments

The authors would like to thank the Swiss Light Source for crystallographic data collection and Tim Sharpe from the Biozentrum Biophysics Facility, Stella Stefanova from the Biozentrum FACS Core Facility, Oliver Biehlmaier from the Biozentrum Imaging Core Facility and Thomas Müntener from the Swiss High-field NMR Facility for experimental support. This study was supported by the Swiss National Science Foundation project grant 310030_189253 to U.J. and by the Swiss National Science Foundation NCCR grant 51NF40_180541 to U.J. We declare no competing financial interests.

---

**Figure S1.**
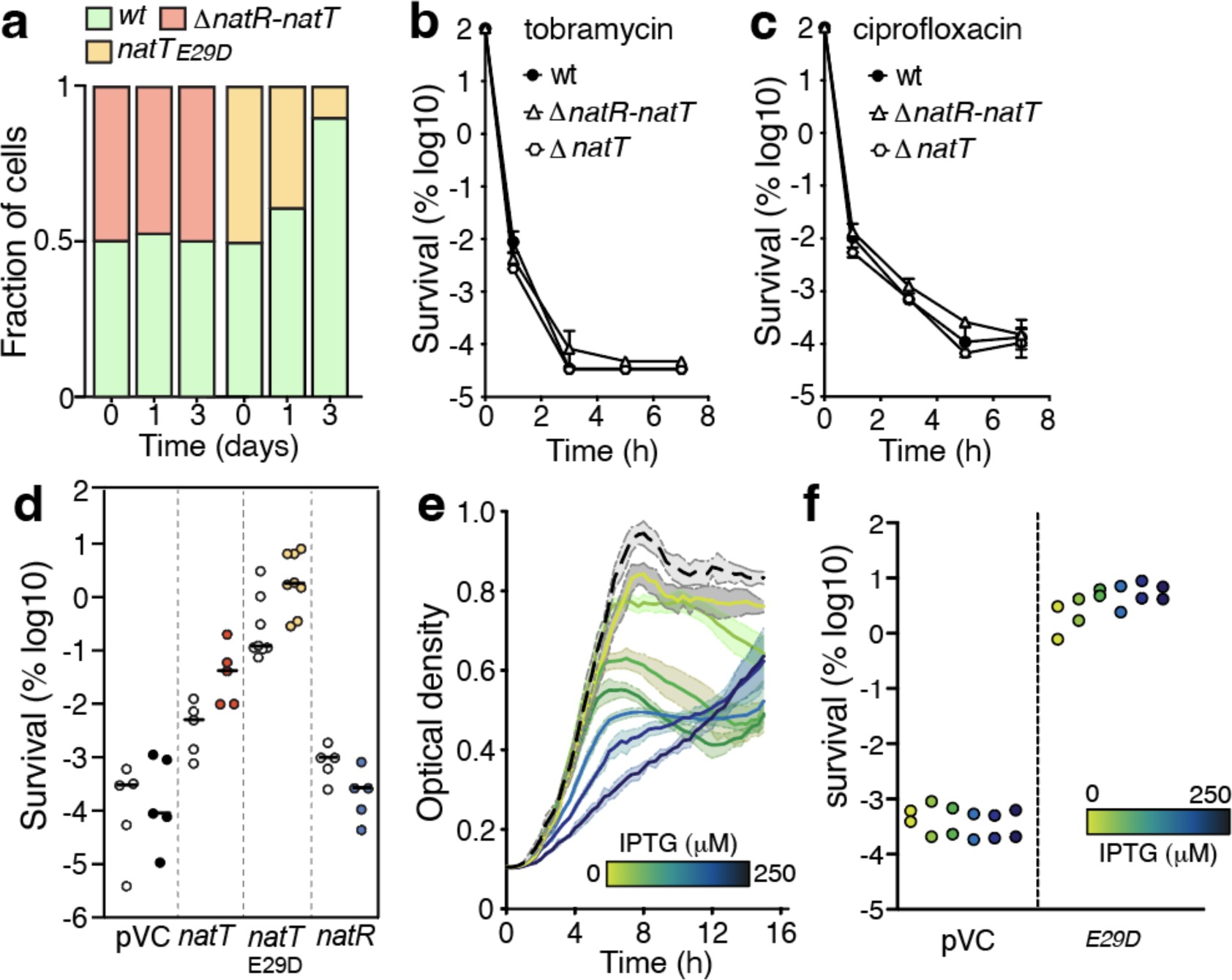
The NatT toxin confers drug tolerance to *P. aeruginosa*. **a)** Reduced fitness of a *P. aeruginosa natTE29D* mutant. *P. aeruginosa* strains expressing different fluorophores were mixed 1:1 and subjected to consecutive cycles of growth and re-dilution. Subpopulations were analyzed by FACS at indicated time intervals. **b,c)** Survival of Δ*natT* and Δ*natR-*Δ*natT* mutants during treatment with tobramycin (8 μg/ml) (**b**) or ciprofloxacin (2.5 μg/ml) (**c**) (average ± SEM, n=3). **d)** Expression of *natT*, but not *natR* provides drug tolerance. Cultures of *P. aeruginosa* containing plasmids with IPTG-inducible *natT* or *natR* alleles were grown in LB medium without (empty boxes) or with 250 μM IPTG (filled boxes). Fractions of surviving cells were determined after three hours of treatment with ciprofloxacin (2.5 μg/ml) (n≥5; lines mark medians) (pVC=control plasmid). **e)** Expression of *natTE29D* gradually limits *P. aeruginosa* growth. Cultures of a Δ*natT* mutant containing a control plasmid (stippled line) or a plasmid expressing *natTE29D* from an inducible promoter, were grown in LB with increasing concentrations of IPTG as indicated (average ± SEM, n>3). **f)** Expression of *natTE29D* increases *P. aeruginosa* tolerance. *P. aeruginosa* cultures containing a plasmid with an IPTG-inducible *natTE29D* were grown as in (**e**) and survival was determined after three hours of treatment with tobramycin (16 μg/ml) (average ± SEM).

**Figure S2.**
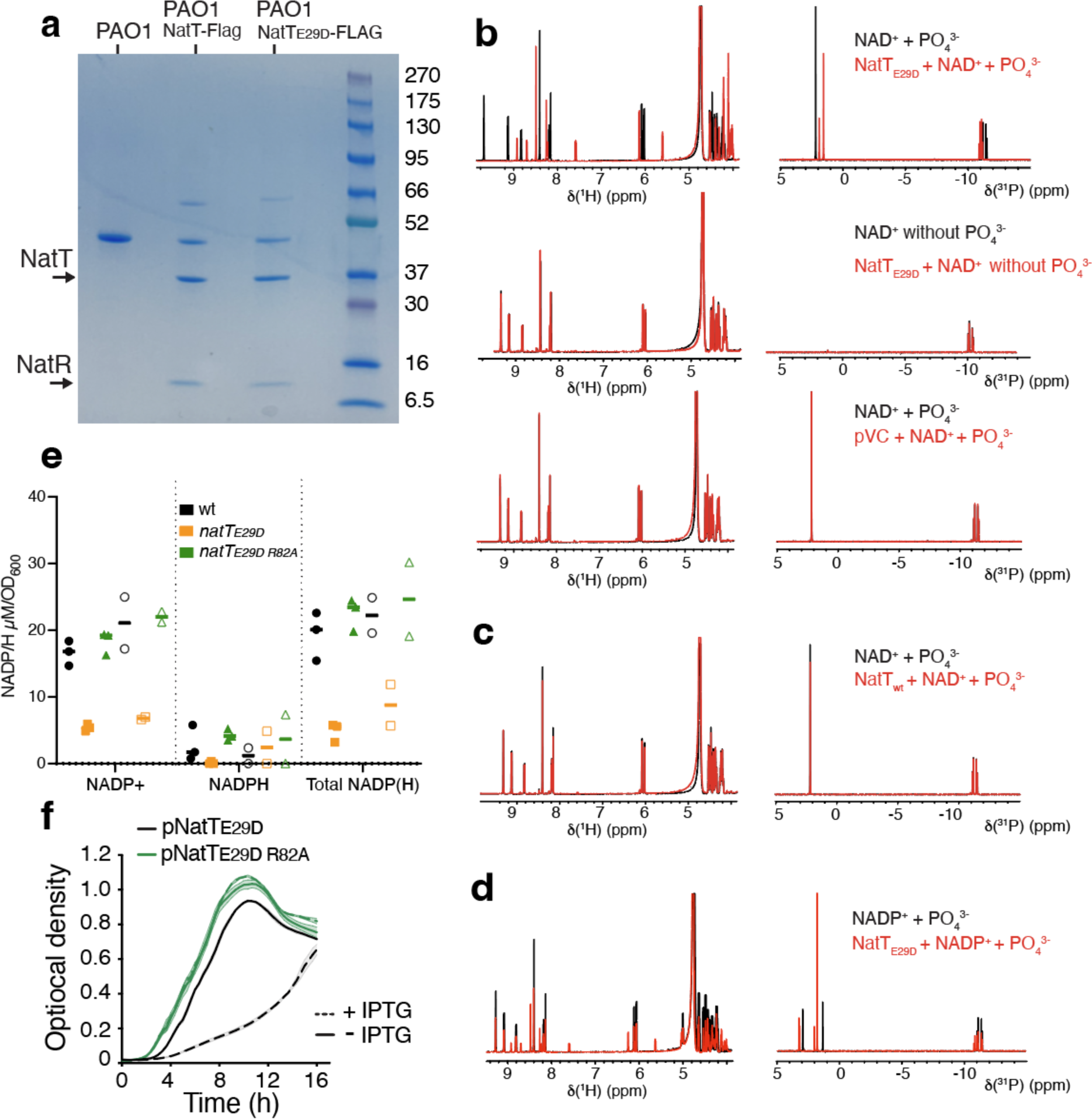
NatT is a NAD-dependent phosphorylase. **a)** Purification of NatR-NatT complex from *P. aeruginosa*. NatR and NatT proteins were copurified from a *P. aeruginosa* Δ*natT* mutants harboring plasmids containing FLAG-tagged *natT* alleles by affinity chromatography using anti-FLAG beads. **b)** Degradation of NAD^+^ by NatTE29D is phosphate-dependent. Purified NatR-NatTE29D complex was incubated with NAD^+^with and without phosphate, and the reaction was analyzed by 1D ^1^H and ^31^P NMR spectra. **c)** Purified NatR-NatT wild-type does not degrade NAD^+^. **d)** Purified NatR-NatTE29D degrades NADP^+^ in a phosphate-dependent reaction. **e)** NatT depletes cellular NADP pools of *P. aeruginosa*. NADP^+^ and NADPH concentrations were determined in cultures of *P. aeruginosa* wild type (wt) and in strains harboring plasmids expressing *natTE29D* or *natTE29D R82A* growing exponentially (filled bars) or from stationary phase (open bars) (average ± SEM, n=2). **f)** The active site residue R82 is required for NatT-mediated toxicity. *P. aeruginosa* with plasmids containing IPTG-inducible copies of *natTE29D* (black line) or *natTE29D R82A* (green line) were grown in LB with (dotted lines) or without IPTG (solid lines) (average ± SEM, n=3).

**Figure S3.**
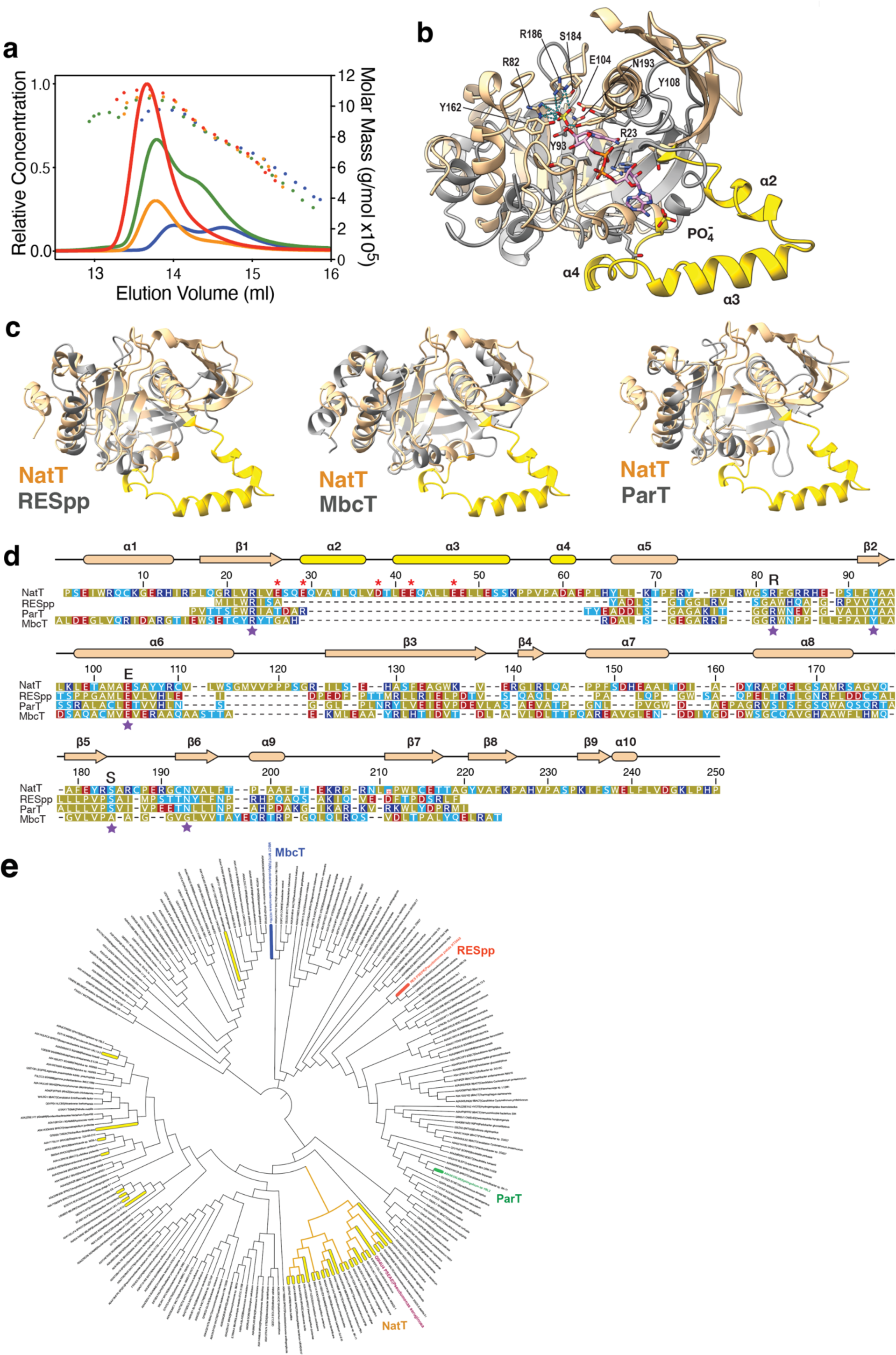
Interaction of NatR and NatT mediates toxin activation. **a)** SEC-MALS analysis of NatR-NatT and NatR-NatTE29D complexes reveal different oligomeric structures. Experiments were performed with 7 µM (red) and 2 µM (orange) NatR-NatT and with 7 µM (green) and 2 µM (blue) NatR-NatTE29D. **b)** Structural homology between NatT and diphtheria toxin indicates analogous NAD^+^ binding modes. NatT is shown in light brown and yellow (Flap), diphtheria toxin (1tox) is in gray and its active site NAD^+^ is in pink. **c)** Superposition of *P. aeruginosa* NatT (light brown) with Flap (yellow) and RES domain proteins RESpp^20^, MbcT^12^ and ParT^14^ (gray). **d)** Structure-guided sequence alignment of NatT and RES domain proteins shown in (**b**). Predicted active site residues are marked with purple stars. Charged residues of the Flap involved in interaction with NatR’ are marked with red asterisks. **e)** Phylogenetic tree of bacterial RES domain proteins with NatT, ParT, RESpp and MbcT highlighted. Branches of RES domains containing a Flap region are highlighted in yellow.

**Figure S4.**
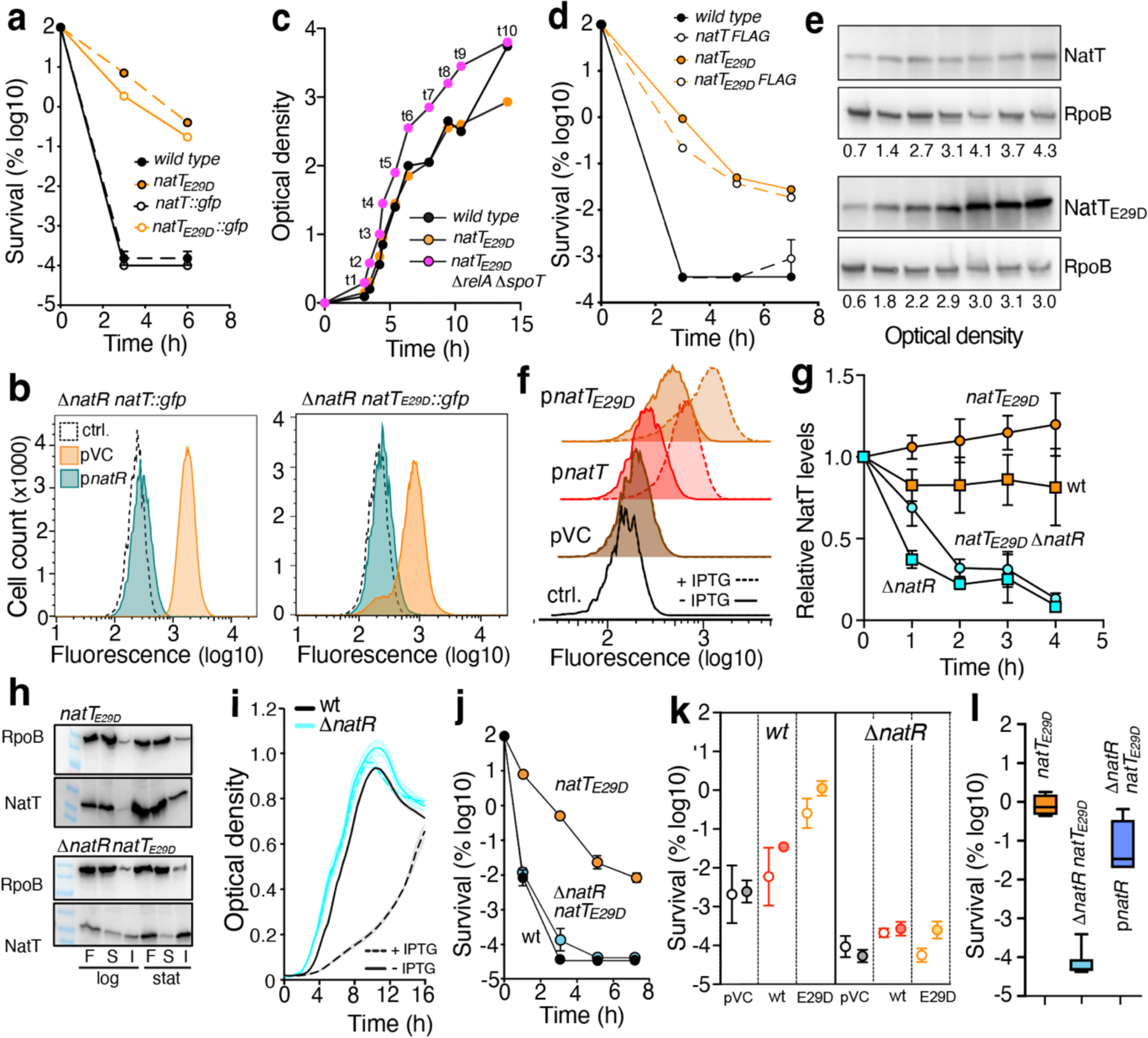
NatR is an antiand a co-toxin of NatT. **a)** Introduction of a chromosomal *natT*::*gfp* reporter does not affect *P. aeruginosa* drug tolerance. Survival of *P. aeruginosa* was scored during exposure to tobramycin (16 μg/ml) (average ± SEM, n>3). **b)** NatR is a repressor of *natT* transcription. Transcription of *natT* was determined by FACS in 11.*natR* mutants with a *gfp* reporter downstream of *natT*. Strains contained a control plasmid (orange) or a plasmid expressing *natR* (green). Control strains lacking the *gfp* reporter are indicated by a dotted black line. **c)** Growth of *P. aeruginosa* wild type, *natTE29D* and *natT*E29D *ΔrelA ΔspoT* in LB. Samples from different time points (t1-t10) were used for FACS analysis in Fig. 4e. **d)** Drug tolerance is not affected by an engineered chromosomal *natT-FLAG* allele (average ± SEM, n=3). **e)** NatT protein levels increase in *P. aeruginosa natTE29D* upon entry into stationary phase. Levels of NatT and NatTE29D were determined during growth (see: **c**) by immunoblot analysis using anti-FLAG antibodies and anti-RpoB antibodies as control. **f)** Ectopic expression of *natT* leads to derepression of *natR-natT* transcription. Transcription of *natR-natT* was determined in *P. aeruginosa* harboring a chromosomal *natT*::*gfp* reporter and plasmids with IPTG-inducible *natT* alleles as indicated (pVC=control plasmid). **g)** NatT is degraded in strains lacking NatR. Concentrations of NatT were determined by immunoblot analysis after treating *P. aeruginosa* cultures with chloramphenicol and plotted as relative values of the initial concentration (0 hrs) (average ± SEM, n≥3). **h)** NatT is insoluble in strains lacking NatR. NatT protein was quantified by immunoblot analysis of fractions harvested from different *P. aeruginosa* strains and in different growth phases as indicated. F: full lysate; S: soluble fraction; I: insoluble fraction. **i)** NatT-mediated toxicity is abolished in a Δ*natR* mutant. Growth of *P. aeruginosa* wild type and Δ*natR* mutant carrying a plasmid with an IPTG-inducible *natTE29D* allele was recorded in LB medium with or without IPTG as indicated (average ± SEM, n=3). **j)** NatR is required for NatT-mediated drug tolerance. Survival of *P. aeruginosa* wild type and mutants indicated during exposure to tobramycin (average ± SEM, n>3). **k)** NatR is required for NatT-mediated drug tolerance. Survival of *P. aeruginosa* wild-type and *ΔnatR* mutant carrying plasmids with IPTG-inducible copies of *natT* or *natTE29D* was determined after treatment with tobramycin (16 μg/ml). Cultures were grown with (filled circles) or without IPTG (empty circles) (average ± SEM, n=3). **l)** Ectopic expression of *natR* restores drug tolerance of a Δ*natR natTE29D* mutant. Survival to tobramycin is shown for strains indicated. Plasmid p*natR* harbors an IPTG-inducible copy of *natR* (average ± SEM, n=3).

**Figure S5.**
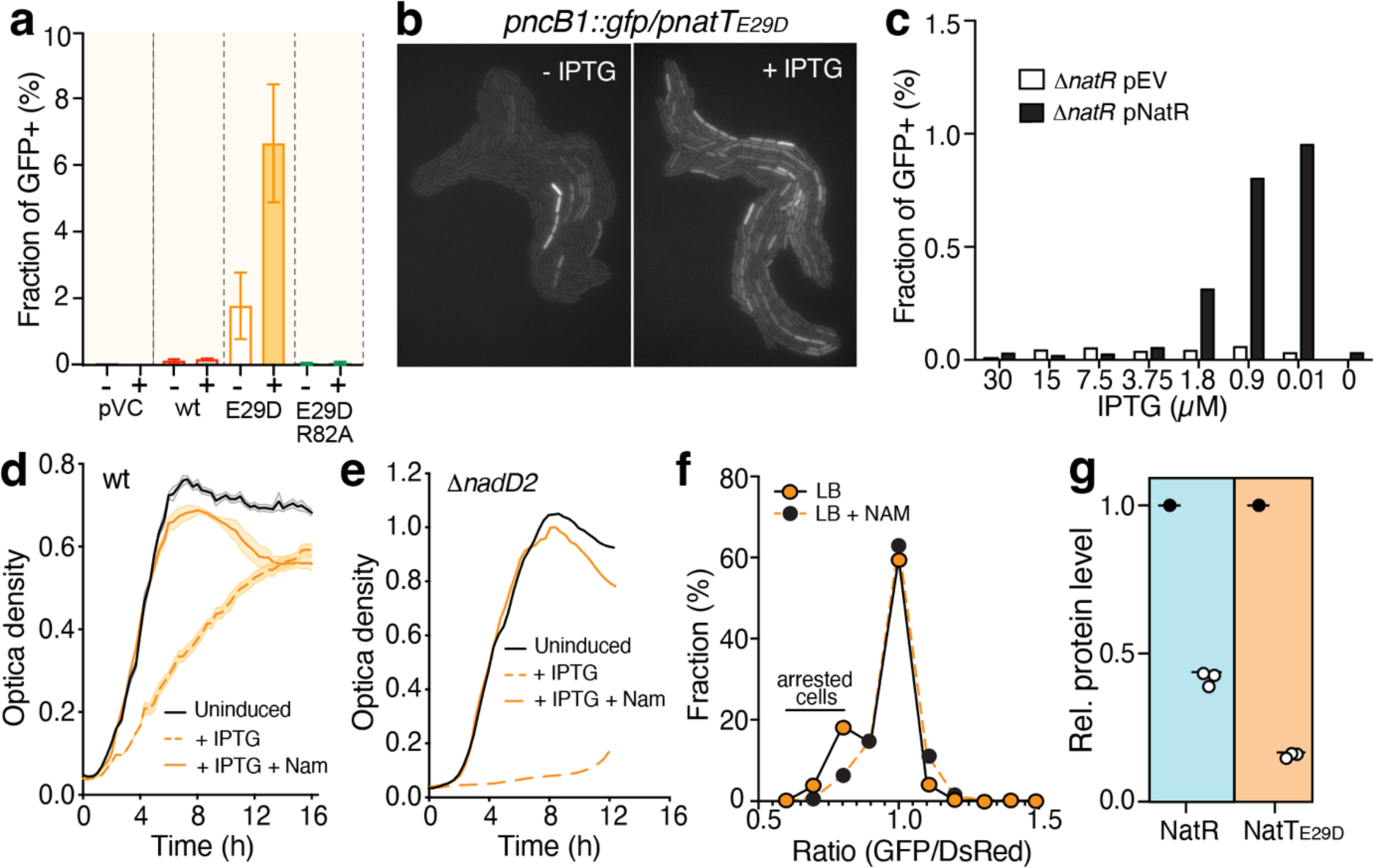
The NAD salvage pathway neutralizes NatT toxin activity and abolishes drug tolerance. **a)** Ectopic expression of *natT* mediates NAD^+^ salvage pathway induction. Fractions of cells inducing salvage pathway genes were determined in *P. aeruginosa pncB1*::*gfp* reporter strains harboring plasmids expressing different *natT* alleles from an IPTG-inducible promoter. Cultures were grown with or without IPTG (average ± SEM, n=3) (pVC=control plasmid). **b)** Ectopic expression *natTE29D* induces salvage pathway genes. Representative microscopy images of a *pncB1*::*gfp* reporter strain expressing *natTE29D* from a plasmid with or without IPTG. **c)** Limiting *natR* expression induces salvage pathway genes. *P. aeruginosa* Δ*nrtR pncB1*::*gfp* carrying a plasmid with an IPTG-inducible *natR* was analyzed at different IPTG concentrations as indicated. Fractions of cells with derepressed salvage pathway were scored as a function of *natR* expression. **d,e)** NAM neutralizes NatTE29D-mediated growth defect in *P. aeruginosa* wild type (**d**) or Δ*nadD2* salvage pathway mutant (**e**). Strains harboring a plasmid with an IPTG-inducible *natTE29D* allele were grown with (orange) or without IPTG (black) and with (solid lines) or without NAM (20mM) (dotted lines). **f)** NAM overrides NatT-mediated growth arrest. Cultures of *P. aeruginosa* wild type and *natTE29D* mutant constitutively expressing TIMER were grown in LB with or without NAM for 3 hrs before analyzing populations by FACS. Fractions of slow growing or arrested cells were calculated using GFP/DsRed ratios of individual cells. **g)** NAM limits NatR and NatT protein levels. A *P. aeruginosa natRE29D* mutant was grown in LB with (open circles) or without NAM (closed circles), followed by the analysis of relative levels of NatR and NatT by mass-spectrometry.

**Figure S6.**
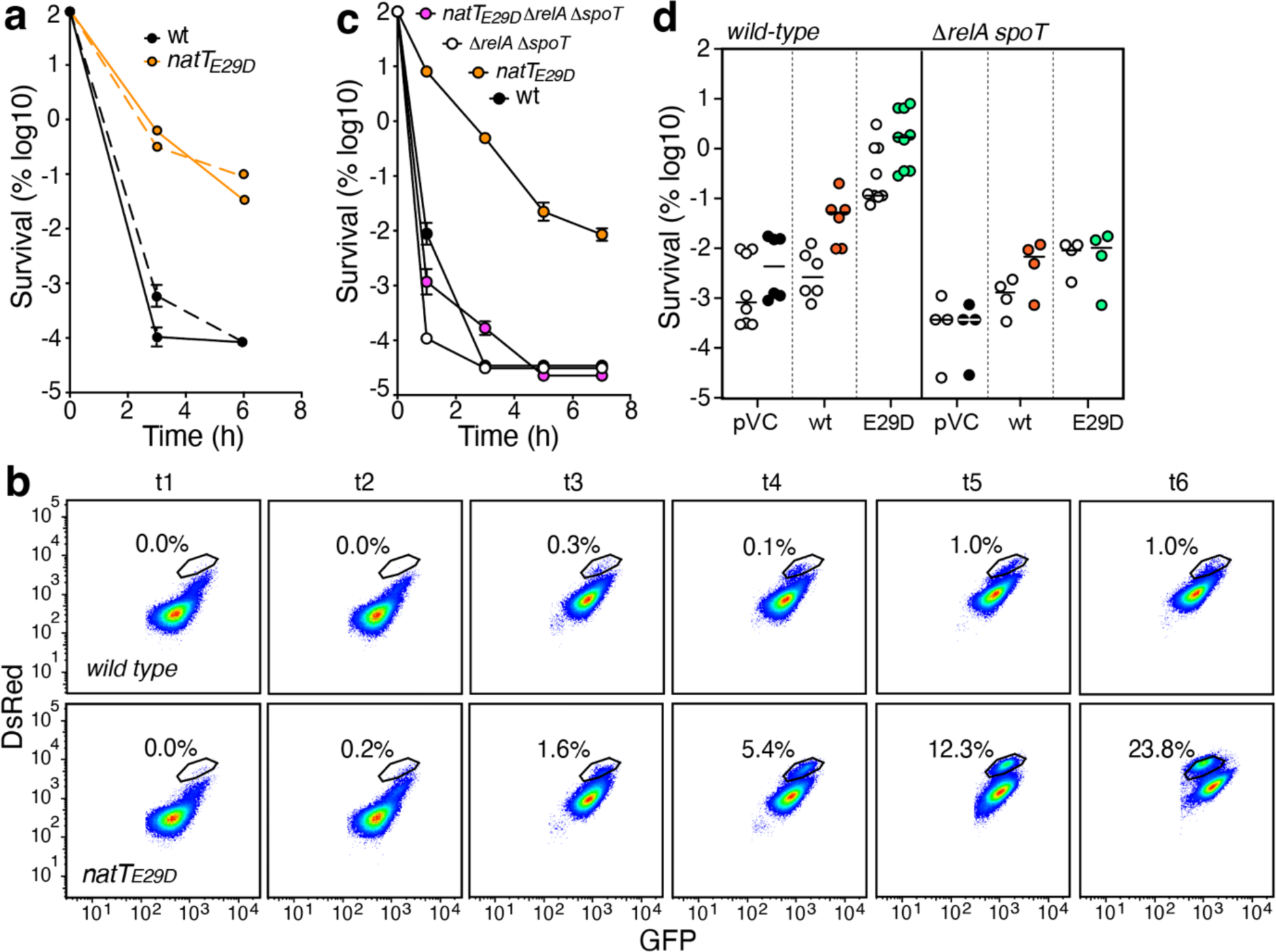
NatT activation generates subpopulations with extended lag phase during outgrowth. **a)** Expression of TIMER from the chromosomal *attb* locus does not affect *P. aeruginosa* drug tolerance. Cell survival during tobramycin treatment was determined for isogenic strains with (dotted line) or without (solid line) TIMER. **b)** The *natTE29D* allele induces growth arrest in a subpopulation of *P. aeruginosa* cells. Samples of *P. aeruginosa* wild type and *natTE29D* mutant cultures expressing TIMER were harvested at different time points (see: Fig. 6a), diluted into fresh medium for three hours, and analyzed by FACS. Fractions of slow-growing cells (black gate) are indicated. **c,d)** (p)ppGpp is required for NatT-mediated drug tolerance. **c)** Cultures of *P. aeruginosa* strains indicated were treated with tobramycin and survival was determined over time. **d)** Survival of *P. aeruginosa* wild-type and Δ*relA* Δ*spoT* mutant carrying plasmids with IPTG-inducible *natT* (wt) or *natTE29D* (E29D) alleles was determined after three hours of treatment with tobramycin. Cultures were grown with (filled circles) or without IPTG (open circles). Solid lines mark median values.

## Materials & Methods

### Bacterial strains and culture conditions

Strains used in this study are listed in Table S2. Unless otherwise stated, *P. aeruginosa* and all *E. coli* strains were grown at 37°C in Luria Bertani (LB) medium under shaking at 170 rpm or on plates containing 1.3% agar where appropriate. Media *P. aeruginosa* were supplemented with 30 µg/ml gentamycin (*E. coli* 20 µg/ml) or 100 µg/ml tetracycline (*E. coli* 12.5 µg/ml). Stocks of 1M isopropyl-β-D-thiogalactopyranoside (IPTG), 1M nicotinamide (NAM), 10 mg/ml tobramycin, 1 mg/ml Ciprofloxacin were prepared in water. Stocks of 1M cumate and 30 mg/ml Chloramphenicol were prepared in 100% ethanol.

### Plasmids and oligonucleotides

Plasmid and primers used in this study are listed in Table S3 and Table S4.

### Molecular biology procedures

Cloning was carried out as previously described^58^. DNA fragments were amplified by PCR using Phusion polymerase. Vectors were cut by restriction and dephosphorylated using calf intestinal alkaline phosphatase. Vectors and inserts were gel purified and ligated at 16°C overnight or room temperature for 10 minutes using T4 DNA Ligase. After ligase deactivation at 65°C for 10 min, chemically competent *E. coli* DH5a cells were transformed by heat shock at 42°C for 45 seconds, followed by phenotypic expression and plating on LB agar containing the appropriate antibiotic. Constructs were validated by sequencing. Details for each construct are provided in Table S2. Primers used in this study are listed in Table S4.

### Chromosomal deletion by homologous recombination

All chromosomal deletions and allelic replacements were engineered via two-step allelic exchange using pEX18-based vectors^59^. For the pEX18Tc-based (Tet^R^) deletion constructs flanking regions (c.a. 700bp) of the target gene were cloned. For pEX18Tc-based constructs to generate chromosomal *gfp* reporter strains, the eGFP gene was cloned between regions flanking the target gene. *Pseudomonas aeruginosa* was transformed by electroporation, resistant colonies selected by picking onto selective plates and then transferred to plates containing 8% sucrose to select for double cross-over. Mutants were validated by colony PCR (deletions, insertions) or by sequencing (allelic exchange). Plasmids used are listed in Table S3.

### Site-Directed Mutagenesis

Individual alleles *of natT* were constructed using the Quick-Change mutagenesis protocol (Stratagene) with Phusion DNA polymerase (NEB) and the respective oligonucleotides according to the manufacturer’s instructions (GE Healthcare). After amplifying the vectors using the mutagenesis primers listed in Table S3, target vectors were digested for 1 hr at 37°C using *Dpn*I restriction endonuclease. Point mutations were verified by DNA sequencing.

### Ectopic expression of *natT* alleles in *P. aeruginosa*

To express different alleles of *natT* or *natR* in *P. aeruginosa*, strains carrying the empty plasmid pME6032 or pME6032 harboring *natT* or *natR* copies were grown ON in LB supplemented with Tet 100 μg/ml and then diluted to an OD 0.05 with fresh LB supplemented with different concentrations of IPTG (0 - 250 μM) and grown in microtiter plates in an Epoch-2 reader (Agilent Technologies). 20 mM NAM was supplemented where indicated.

### Antibiotic killing assays

To quantify *P. aeruginosa* survival during antibiotic treatment, cultures were inoculated from a single colony in LB or in LB supplemented with NAM, grown to different ODs as indicated and then diluted back with fresh LB medium containing antibiotics to a standard OD of 0.12. Unless otherwise stated, tobramycin and ciprofloxacin were used at 16 µg/ml and 2.5 µg/ml, respectively. Culture aliquots were collected at regular time intervals, diluted and plated on LB plates to determine survival frequencies. *P. aeruginosa* strains carrying derivatives of plasmid pME6032 carrying different *natR* or *natT* alleles were grown in LB supplemented with 100 μg/ml tetracycline, diluted to OD 0.1 in LB Tet 100 μg/ml with or without 250 μM IPTG and grown for four hours to allow the expression of the protein. Bacteria were then diluted 1:2 in LB supplemented with 16 µg/ml tobramycin. Culture aliquots were collected at regular time intervals, diluted and plated on LB plates to determine survival frequencies.

### Determination of minimal inhibitory concentrations (MIC)

MIC values were determined by adapting existing protocols^60,61^. Briefly, an overnight culture was diluted in LB to reach a density of 10^6^ CFU/ml, and grown in the presence of increasing concentrations of antibiotics for 16–20 hrs at 37°C with shaking. After incubation, the OD_600_ was determined and MIC values were determined as the lowest antibiotic concentration where no growth was observed.

### Competition experiments

Two different strains constitutively expressing eGFP and mCherry, respectively, were mixed 1:1 in 5 ml LB medium and cultures were grown to stationary phase ON. Cultures were then diluted back 1:100 into fresh LB growth cycles were repeated for 3 days. Culture aliquots were removed daily, diluted in PBS and used to determine their composition by flow cytometry.

### Flow cytometry

To determine subpopulations with different lag periods during outgrowth, strains constitutively expressing TIMER^bac34^ from the chromosome were inoculated in LB and grown into stationary phase. Bacteria were diluted with fresh LB to an OD of 0.12, incubated for 3 hours before the fraction of green and red cells was determined by flow cytometry. To determine *natT* expression or *pcnB1* expression in individual cells, cultures of strains harboring a chromosomal *natT::gfp* or *pcnB1::gfp* fusion were inoculated in LB medium. At the indicated time points culture aliquots were removed, diluted in PBS and analyzed. Measurements were performed on a BD LSR Fortessa 4 Laser and fluorescence determined with the following channel, Ex488_LP495_BP514/30-H for eGFP and Ex561_LP600_BP610/20-H for mCherry.

### FACS sorting

Cells expressing TIMER^bac34^ were inoculated in LB and grown into stationary phase, before being diluted back to an OD of 0.12 with fresh LB or LB with 20 mM NAM, and grown for three hours. Cell cultures were sorted according to their GFP/mCherry ratio using an BD FACSAria IIIu (BD Biosciences) with scatter and fluorescence channels (green, Ex488_LP495_BP514/30-H; mCherry, Ex561_LP600_BP610/20-H). Triplicates of sorted cells (4×10^5^) were inoculated in 1 ml LB containing tobramycin (10 mg/ml) and incubated for different time intervals before aliquots were removed plated on LB plates to determine survival rates. Aliquots of sorted populations were directly plated on LB plates to determine the number of viable bacteria after sorting.

### Fluorescence microscopy

Overnight cultures were diluted 1:100 in fresh media and transferred to 1% agarose pads containing LB medium and IPTG where indicated. Time-lapses were recorded on a DeltaVision microscope (Applied Precision) equipped with a 100x oil immersion objective and an environmental chamber maintained at 35°C. Images were recorded using a CoolSnap HQ2 camera and processed using Softworx software (Applied Precision).

### Immunoblot analysis

Strains expressing FLAG-tagged versions of NatT were inoculated in LB medium or in LB supplemented with 20 mM NAM and harvested in different growth phases. Aliquots were sampled at different times points, pelleted by centrifugation and snap frozen in liquid nitrogen. Samples were dissolved in SDS sample buffer containing 1% betamercaptoethanol and 2% SDS, boiled for 5 min and separated by SDS-PAGE (15%). Proteins were semi-dry transferred to a nitrocellulose membrane (Amersham Protan 0.2μm), membranes were blocked overnight in PMT (PBS, 0.1% v/v Tween 20, 5% w/v dry milk) and stained with polyclonal anti-FLAG M2 antibodies (1:10,000) (Sigma) and with polyclonal swine anti-mouse-HRP antibodies (1:1,000, Dako, Denmark). Protein bands were stained with ECL reagents and chemiluminescence was detected with a LAS4000 imager using ImageQuant LAS4000 version 1.3.

### Metabolome analysis

Cells were pelleted and snap frozen in liquid nitrogen. Metabolites were extracted twice with hot (> 70°C) 60% ethanol. Extracts were analyzed by flow injection – time of flight mass spectrometry on an Agilent 6550 QTOF instrument operated in the negative mode, as described previously^62^.

### Quantification of NAD and NADP

The concentrations of NAD+/NADH and NADP+/NADPH were determined using Assay Kits from Fluorometric following the manufacturer’s instructions. Overnight cultures of *P. aeruginosa* were diluted 1:100 in LB, grown to different optical densities before aliquots were removed, washed twice in PBS and resuspended in the lysis buffer provided by the manufacturer.

### Protein expression and purification

Overnight cultures of *E. coli* BL21 (DE3) (Thermo Scientific) carrying a pRSF-Duet-1 plasmid with *natR* and a His-tagged copy of *natT* were diluted 100x in 4L of fresh LB medium supplemented with kanamycin at 100 µg/ml. Expression was induced by the addition of 500 µM IPTG at an O.D of 0.8 for 16 hours at 18 °C. Cells were harvested by centrifugation at 7000 xg for 15 minutes and pellets frozen at -80°C. Before lysis, the cell pellet was resuspended in 50 ml of a solution containing 50 mM Bicine pH 8.5, 300 mM KCl, 10 mM TCEP, 0.5 mg/ml lysozyme and protease inhibitor (cOmplete, Mini, EDTA-free, Roche). Lysis was performed by 3 passages through a cell disruptor, and the insoluble fraction was separated from soluble protein by ultracentrifugation for 30 min at 135,000 xg. The supernatant was loaded on a HisTrap column HP 5 ml (Cytiva) equilibrated with 50 mM Bicine pH 8.5, 300 mM KCl, 10 mM TCEP, and 20 mM Imidazole. Proteins were eluted by an imidazole gradient (20 mM to 500 mM) in 15 column volumes. Fractions were analyzed SDS-PAGE gel to assess protein quality and purity. Protein fractions were pooled, concentrated and submitted to a size exclusion chromatography S200 16/60 (Cytiva) at a flow of 0.5 ml/min in 50 mM Bicine pH 8.5, 50 mM KCl, 10 mM TCEP. Protein quality was again assessed by SDS-PAGE and fractions corresponding to the NatR-NatT complex were concentrated to 3-5 mg/ml, aliquoted and flash frozen with liquid nitrogen and stored at -80°C. The same protocol was applied for NatT_E29D_ mutant protein complexes.

NatT samples for enzymatic assays were obtained directly from *P. aeruginosa* strains carrying plasmid pME6032 expressing FLAG-tagged versions of *natT* or *natT_E29D_*. *P. aeruginosa* cultures (500 ml) were induced with IPTG (500 µM) for 3 hours at 37°C at OD 0.8. Cells were harvested by centrifugation at 7000 xg for 15 minutes and were lysed in 10 ml PBS supplemented with protease inhibitor (cOmplete, Mini, EDTA-free, Roche). NatT was purified by immunoprecipitation using Anti-FLAG magnetic beads (Merk) and eluted with FLAG peptide following the manufacturer’s instructions. Protein quality was assessed by SDS-PAGE and identical protocols were applied to NatT and NatT_E29D._

### Mass photometry analysis of NatRT complex and NatRT complex with dsDNA

Mass photometry was carried out on a Refeyn OneMP instrument (Refeyn Ltd.) in microscope coverslips of 24 x 50 mm in size and 1.5 mm thickness (Marienfeld, Cat. No. 0107222). The coverslips were washed with water, 50% isopropanol, and dried using compressed air according to the manufacturer’s instructions. In order to form drops for the measurement, silicone gaskets were used on top of the coverslips (Merck, GBL103250). The instrument was calibrated with molecular weight markers (NativeMark^TM^ Unstained Protein standard, Invitrogen, Cat. No. LC0725). Focus for the measurement was found using 18 µl of buffer without protein (50 mM Bicine pH 8.5, 50 mM KCl). After locking the focus for the measurement, 2 µl of the sample was added to the initial 18 µl of the buffer. Measurements were performed with NatR-NatT complexes at a final concentration of 10 nM. For DNA binding studies, dsDNA corresponding to the *natR* promoter region was added to the measurements with the protein complex at a final concentration of 1 µM. Movies were recorded for 60 seconds using Refeyn Acquire^MP^ software (version 2.5.1) and analyzed with Refeyn Discover^MP^ (version 2.5.0).

### NatR-NatT crystallization and data collection

Se-Met crystals of NatR-NatT complexes were produced by sitting drop vapor diffusion method in a crystallization solution composed of 0.25 M di-Ammonium tartrate and 17.8% PEG 3,350, optimized from the condition n° 86 of the commercial screen NeXtal PEG suit HT (Molecular dimensions). NatR-NatT was crystallized in 0.2 M Sodium sulfate 0.1 M Bis-Tris propane 7.5 20% w/v PEG 3350 from the crystallization screen Pact Premier (Molecular dimensions) at a final protein concentration of 2.5 mg/ml. NatR-NatT_E29D_ crystals were formed in 0.2 M sodium/potassium phosphate pH 7.5, 0.1 M HEPES 7.5, 22.5 % v/v, PEG Smear Medium, 10 % v/v Glycerol from condition G11 of BCS screen (Molecular dimensions) at a final concentration of 2.5 mg/ml. The crystallization plates were set by a Gryphon robot (Art Robbins Instruments) and stored at 20°C. All crystals appeared between 4 to 7 days in drops prepared at 1:1 v/v protein:crystallization solution in a total volume of 0.4 µl.

### Data collection and structure determination

Data were collected at the Swiss Light Source (SLS), Villigen, Switzerland, at beamline X06DA - PXIII using DA + data acquisition software and were processed by XDS and CCP4i2 suite. Se-Met NatR-NatT crystals were diffracted at 3.2 Å, and an initial structure was obtained by single-wavelength anomalous dispersion (SAD) using Crank2 software. Native NatR-NatT crystals were diffracted at 2.3 and 2.4 Å, and the structure was solved by molecular replacement using the preliminary structure obtained from SAD. Model building was carried out on COOT, and refinement was performed using Phenix. Structure figures were prepared in ChimeraX. NatR-NatT and NatR-NatT_E29D_ models were deposited at the Protein Data Bank under the accession codes 8QNL and 8QNQ, respectively. Statistics of the crystallographic data and protein refinement are shown in Table S5.

### NAD+ phosphorylase assay

Purified NatR-NatT_E29D_ complex at a concentration of 40 nM was used in enzymatic assays with 5 mM of NAD^+^ in 25 mM Tris pH 8.0, 300 mM KCl in the presence or absence of 20 mM Potassium phosphate in a total volume of 500 µl. Reactions were started by adding the enzyme to a microcentrifuge tube, and the content was immediately transferred to an NMR tube. For end-point products analysis the reactions were carried out for 16 h and 1D ^1^H, 1D ^31^P and 2D [^31^P,^1^H]-HMBC NMR spectra were recorded. For the determination of the NatT kinetic parameters a series of 1D ^1^H spectra were recorded sequentially (20 minutes per experiment) and the concentration of NAD+ determined by integration of the NMR signals. The NMR experiments were recorded at 298 K on a Bruker Ascend 600 MHz spectrometer running Topspin 3.2 equipped with a cryogenically cooled triple-resonance probe. All data were processed with Bruker TOPSPIN-NMR software (version 3.2, Bruker).

### *Analysis of natT* and *natR* in clinical isolates

Sequences of *natT* and *natR* were obtained from 9594 *Pseudomonas aeruginosa* raw genomic sequences from the Sequence Read Archive database (SRA) using blastn_vdb tool and *natT* from PAO1 as input sequence. A total of 8286 *natT* and 8266 *natR* sequences were identified. Raw reads were translated and SNPs were spotted. Variant *natT* alleles were amplified by colony PCR from a collection of 300 clinical isolates of *P. aeruginosa*. The resulting PCR products were sequenced and cloned into pME6032.

### Ethics statement

Clinical isolates of *P. aeruginosa* used in this study were cultured from patient samples collected for routine microbiological testing at the University Hospital, Basel. Sub-culturing and analysis of bacteria were performed anonymously. No additional procedures were carried out on patients. Cultures were sampled following regular procedures with written informed consent, in agreement with the guidelines of the Ethikkommission beider Basel EKBB.

