## Supplementary tables for "Toxin-mediated depletion of nicotinamide dinucleotides drives persister formation in a human pathogen" (from another version of this preprint: version 1, 2023-09-29)

**Supplementary Table S1:** Polymorphism analysis of *natT* sequences from 8286 *Pseudomonas aeruginosa* genomes.

Genome sequences of *P. aeruginosa* were retrieved from the Sequence Read Archive database (SRA, https://www.ncbi.nlm.nih.gov/sra) and *natT* sequences were identified (N=8286) as reported in the sheet of **Table S1a** of the accompanying Excel file "**Supplementary Table 1**". From left to right, the table lists SRA references of the genomes, sample isolation source, as well as strains and SNPs identified in the corresponding genomes. A pivot table containing all data is shown in sheet **Table S1b** of the Excel. From left to right, the table lists isolation categories of samples and, for each category, the number of sequencing datasets, the fraction of samples with a *natT* sequence differing from P. aeruginosa strain PAO1, the number of distinct alleles identified, the relative genetic variability of *natT* as the ratio between the number of distinct alleles and the number of sequenced datasets. Additional columns report the occurrence counts for each allele.

**Supplementary Table S2:** Strains

| Strains | Description | Reference |
|---|---|---|
| PAO1 | WT *P. aeruginosa* | 1 |
| $natT_{E29D}$ | Chromosomal point mutation of the *natT* gene at the original *natT* locus | 2 |
| $natT_{E29D\ R82A}$ | Chromosomal point mutation of the *natT* gene at the original *natT* locus | This study |
| Δ*natT* | Unmarked deletion of *natT* (PA1030) in PAO1 | This study |
| Δ*natR* | Unmarked deletion of *natR* (PA1029) in PAO1 | This study |
| Δ*natRT* | Unmarked deletion of *natR* (PA1029) and *natT* (PA1030) in PAO1 | This study |
| Δ*natR* $natT_{E29D}$ | Unmarked deletion of *natR* (PA1029) in $natT_{E29D}$ background | This study |
| $natRT_{wt}$*::gfp* | Chromosomal fluorescent transcriptional reporter of *natRT* operon in PAO1 | This study |
| Δ*natR* $natRT_{wt}$*::gfp* | Unmarked deletion of *natR* (PA1029) in $natRT_{wt}$*::gfp* background | This study |
| Δ*natT* $natRT_{wt}$*::gfp* | Unmarked deletion of *natT* (PA1030) in $natRT_{wt}$*::gfp* background | This study |
| $natRT_{E29D}$*::gfp* | Chromosomal fluorescent transcriptional reporter of *natRT* operon in $natT_{E29D}$ background | This study |
| Δ*natR* $natT_{E29D}$*::gfp* | Unmarked deletion of *natR* (PA1029) in $natRT_{E29D}$*::gfp* background | This study |
| $natT_{wt}$*-FLAG* | PAO1 with $natT_{wt}$*-FLAG*, with the flag tag fused on the 3' end at the original *natT* locus | This study |
| Δ*natR* $natT_{wt}$*-FLAG* | Unmarked deletion of *natR* (PA1029) in $natT_{wt}$*-FLAG* background | This study |
| $natT_{E29D}$*-FLAG* | $natT_{E29D}$ with $natT_{E29D}$*-FLAG*, with the flag tag fused on the 3' end at the original *natT* locus. | This study |
| Δ*natR* $natT_{E29D}$*-FLAG* | Unmarked deletion of *natR* (PA1029) in $natT_{E29D}$*-FLAG* background | This study |
| *pcnB1::gfp* | Chromosomal fluorescent transcriptional reporter of *pcnB1* (PA4919) in PAO1 | This study |
| $natT_{E29D}$ *pcnB1::gfp* | Chromosomal fluorescent transcriptional reporter of *pcnB1* (PA4919) in $natT_{E29D}$ | This study |
| Δ*natR* $natT_{E29D\ R82A}$ *pcnB1::gfp* | Unmarked deletion of *natR* (PA1029) in $natT_{E29D}$ *pcnB1::gfp* background | This study |
| $natT_{E29D\ R82A}$ *pcnB1::gfp* | Chromosomal fluorescent transcriptional reporter of *pcnB1* (PA4919) in $natT_{E29D\ R82A}$ background | This study |
| *ΔrelA spoT pcnB1::gfp* | Chromosomal fluorescent transcriptional reporter of *pcnB1* (PA4919) in *ΔrelA spoT* background | This study |
| $natT_{E29D}$ *ΔrelA spoT pcnB1::gfp* | Chromosomal fluorescent transcriptional reporter of *pcnB1* (PA4919) in $natT_{E29D}$ *ΔrelA spoT* background | This study |
| *pcnB1::gfp ΔnrtR* | Unmarked deletion of *nrtR* (PA4916) in *pcnB1::gfp* background | This study |
| $natT_{E29D}$ *pcnB1::gfp ΔnrtR* | Unmarked deletion of *nrtR* (PA4916) in $natT_{E29D}$ *pcnB1::gfp* background | This study |
| $natT_{E29D}$ *ΔnrtR* | Unmarked deletion of *nrtR* (PA4916) in $natT_{E29D}$ background | This study |

| | | |
|---|---|---|
| *natT*$_{E29D}$ *ΔnadD2* | Unmarked deletion of *nadD2* (PA4917) in *natT*$_{E29D}$ background | This study |
| *ΔnadD2* | Unmarked deletion of *nadD2* (PA4917) in PAO1 background | This study |
| *ΔrelA spoT* | Unmarked deletion of *relA* (PA0934) and *spoT* (PA5338) in PAO1 background | This study |
| *natT*$_{E29D}$ *ΔrelA spoT* | Unmarked deletion of *relA* (PA0934) and *spoT* (PA5338) in *natT*$_{E29D}$ background | This study |
| *natT*$_{E29D}$*-FLAG ΔrelA spoT* | Unmarked deletion of *relA* (PA0934) and *spoT* (PA5338) in *natT*$_{E29D}$*-FLAG* background | This study |
| *natT*$_{E29D}$*::gfp ΔrelA spoT* | Unmarked deletion of *relA* (PA0934) and *spoT* (PA5338) in *natT*$_{E29D}$*::gfp* background | This study |
| *attb::*TIMER$^{bac}$ | Chromosomal copy of of TIMER$^{bac}$ at *attb* locus | This study |
| *natT*$_{E29D}$ *attb::*TIMER$^{bac}$ | *natT*$_{E29D}$ with a chromosomal copy of TIMER$^{bac}$ at *attb* locus | This study |
| *Tn::nadD* | *Tn::PA1004* | Manoil lab |
| *Tn::nadB* | *Tn::PA0761* | Manoil lab |
| *Tn::nadC* | *Tn::PA4524* | Manoil lab |
| *Tn::pcnA* | *Tn::PA4918* | Manoil lab |
| *Tn::pcnB1* | *Tn::PA4919* | Manoil lab |
| *Tn::nadD2* | *Tn::PA4917* | Manoil lab |
| *Tn::nadE* | *Tn::PA4920* | Manoil lab |
| *Tn::pcnB2* | *Tn::PA4376 (homologue to PA4919)* | Manoil lab |
| *Tn::nadD1* | *Tn::PA4006 (homologue to PA4917)* | Manoil lab |

**Supplementary Table S3:** Plasmids

| Constructs | Description | Construction | Reference |
|---|---|---|---|
| miniCTX::*pX2-gfp* | mini-CTX for *gfp* integration into *attB* locus under control of the pX2 constitutive promoter ($Tet^R$) | - | Dirk Bumann, |
| miniCTX::*pX2-mCherry* | mini-CTX for *mCherry* integration into *attB* locus under control of the pX2 constitutive promoter ($Tet^R$) | - | Dirk Bumann, |
| miniCTX::*pX2-TIMER*$^{bac}$ | mini-CTX for *TIMER*$^{bac}$ integration into *attB* locus under control of the pX2 constitutive promoter ($Tet^R$) | - | Dirk Bumann, |
| pME6032 | $Tet^R$, $P_K$, 9.8 kb pVS1 derived shuttle vector | - | 3 |
| pME6032::$natT_{wt}$ | pME6032 with $natT_{wt}$ under the control of IPTG promoter | *natT* (PA1030) was amplified from PAO1 genomic DNA with primers A and B as *Eco*RI-*Kpn*I fragment | This study |
| pME6032::$natT_{wt\text{-}ATG}$ | pME6032 with $natT_{wt\text{-}ATG}$ under the control of *lac* promoter | *natT* (PA1030) was amplified from PAO1 genomic DNA with primers C and B as *Eco*RI-*Kpn*I fragment | This study |
| pME6032::$natT_{E29D}$ | pME6032 with $natT_{E29D}$ under the control of *lac* promoter | *natT* (PA1030) was amplified from $natT_{E29D}$ genomic DNA with primers A and B as *Eco*RI-*Kpn*I fragment | This study |
| pME6032::$natT_{E29D\ R82A}$ | pME6032 with $natT_{E29D\ R82A}$ under the control of *lac* promoter | *natT* (PA1030) was amplified from $natT_{E29D\ R82A}$ genomic DNA with primers A and B as *Eco*RI-*Kpn*I fragment | This study |
| pME6032::*natR* | pME6032 with *natR* under the control of *lac* promoter | *natR* (PA1029) was amplified from PAO1 genomic DNA with primers D and E as *Eco*RI-*Kpn*I fragment | This study |
| pME6032::$natT_{V117A}$ | pME6032 with $natT_{V117A}$ under the control of *lac* promoter | *natT* (PA1030) was amplified from CI_1125 genomic DNA with primers A and B as *Eco*RI-*Kpn*I fragment | This study |
| pME6032::$natT_{I125T}$ | pME6032 with $\mathrm{natT}_{\mathrm{I125T}}$ under the control of *lac* promoter | *natT* (PA1030) was amplified from CI_1186 genomic DNA with primers A and B as *Eco*RI-*Kpn*I fragment | This study |
| pME6032::$\mathrm{natT}_{\mathrm{S170T}}$ | pME6032 with $natT_{S170T}$ under the control of *lac* promoter | *natT* (PA1030) was amplified from CI_1129 genomic DNA with primers A and B as *Eco*RI-*Kpn*I fragment | This study |
| pME6032::$natT_{I14V\ V117A}$ | pME6032 with $natT_{I14V\ V117A}$ under the control of *lac* promoter | *natT* (PA1030) was amplified from CI_1107 genomic DNA with primers A and B as *Eco*RI-*Kpn*I fragment | This study |
| pME6032::$natT_{V117A\ D151G}$ | pME6032 with $natT_{V117A\ D151G}$ under the control of *lac* promoter | *natT* (PA1030) was amplified from CI_1115 genomic DNA with primers A and B as *Eco*RI-*Kpn*I fragment | This study |
| pME6032::$natT_{I142M\ S170T}$ | pME6032 with $natT_{I142M\ S170T}$ under the control of *lac* promoter | *natT* (PA1030) was amplified from CI_1178 genomic DNA with primers A and B as *Eco*RI-*Kpn*I fragment | This study |
| pME6032::$natT_{T100A\ S170T}$ | pME6032 with $natT_{T100A\ S170T}$ under the control of *lac* promoter | *natT* (PA1030) was amplified from CI_1138 genomic DNA with primers A and B as *Eco*RI-*Kpn*I fragment | This study |
| pME6032::$natT_{I14V\ V117A\ P165S}$ | pME6032 with $natT_{I14V\ V117A\ P165S}$ under the control of *lac* promoter | *natT* (PA1030) was amplified from CI_1124 genomic DNA with primers A and B as *Eco*RI-*Kpn*I fragment | This study |
| pME6032::$natT_{I14V\ V117A\ S170T}$ | pME6032 with $natT_{I14V\ V117A\ S170T}$ under the control of *lac* promoter | *natT* (PA1030) was amplified from CI_1128 genomic DNA with primers A and B as *Eco*RI-*Kpn*I fragment | This study |
| pME6032::$natT_{I14V\ V117A\ S232A}$ | pME6032 with $natT_{I14V\ V117A\ S232A}$ under the control of *lac* promoter | *natT* (PA1030) was amplified from PA14 genomic DNA with primers A and B as *Eco*RI-*Kpn*I fragment | This study |
| pME6032::$natT_{I14V\ S105C\ V117A\ P165S\ S232A}$ | pME6032 with $natT_{I14V\ S105C\ V117A\ P165S\ S232A}$ under the control of *lac* promoter | *natT* (PA1030) was amplified from CI_1179 genomic DNA with primers A and B as *Eco*RI-*Kpn*I fragment | This study |

| | | | |
|---|---|---|---|
| pME6032::*natT*$_{I14V\ S105C\ V117A\ E167K\ S170T}$ | pME6032 with *natT*$_{I14V\ S105C\ V117A\ E167K\ S170T}$ under the control of *lac* promoter | *natT* (PA1030) was amplified from CI_1160 genomic DNA with primers A and B as *EcoR*I-*Kpn*I fragment | This study |
| pME6032::*natT*$_{wt}$-*FLAG* | pME6032 with C-terminally 3xFLAG-tag *natT* under the control of *lac* promoter | *natT* (PA1030) was amplified from PAO1 genomic DNA with primers A and F as *EcoR*I-*Kpn*I fragment | This study |
| pME6032::*natT*$_{E29D}$-*FLAG* | pME6032 with C-terminally 3xFLAG-tag *natT*$_{E29D}$ under the control of *lac* promoter | *natT* (PA1030) was amplified from PAO1 genomic DNA with primers A and F as *EcoR*I-*Kpn*I fragment | This study |
| pGn6032 | GmR, pME6032 based vector, Tet cassette replaced with *aacCI* gentamycin resistance cassette | - | 4 |
| pGn6032::*natT*$_{wt}$ | pGn6032 with *natT*$_{wt}$ under the control of *lac* promoter | *natT* (PA1030) was amplified from PAO1 genomic DNA with primers A and B as *EcoR*I-*Kpn*I fragment | This study |
| pGn6032::*natT*$_{E29D}$ | pGn6032 with *natT*$_{E29D}$ under the control of *lac* promoter | *natT* (PA1030) was amplified from *natT*$_{E29D}$ genomic DNA with primers A and B as *EcoR*I-*Kpn*I fragment | This study |
| pQF | Tet$^{R}$, pQF based vector | | 5 |
| pQF::*natR* | pQF with *natR* under the control of cumate-inducible promoter | *natR* (PA1029) was amplified from PAO1 genomic DNA with primers G and H as *Hind*III-*Kpn*I fragment | This study |
| pRSF-Duet-1_His-NatT-NatR | pRSF-Duet-1 (Novagen) with N-terminally His-tag NatT and NatR under the control of *lac* promoter | *natT* (PA1030) was amplified with primers I and J as *BamH*I-*EcoR*I fragment and *natR* (PA1029) was amplified with primers K and E as *EcoRV*-*Kpn*I fragment from PAO1 genomic DNA | This study |
| pRSF-Duet-1_His-NatT$_{E29D}$-NatR | pRSF-Duet-1 (Novagen) with N-terminally His-tag NatT$_{E29D}$ and NatR under the control of *lac* promoter | *natT* (PA1030) was amplified with primers I and J as *BamH*I-*EcoR*I fragment and *natR* (PA1029) was amplified with primers K and E as *EcoRV*-*Kpn*I fragment from PAO1 genomic DNA | This study |
| **Deletion constructs** | | | |
| pEX18-Tc | *oriT*+, *sacB*+, gene replacement vector, Tc$^{R}$ | - | 6 |
| pEX18-Tc-*natT*$_{E29D}$ | pEX18-Tc carrying the G to T in the *natT* allele as *Hind*III-*Xba*I fragment | - | 2 |
| pEX18-Tc-*natT*$_{E29D\ R82A}$ | pEX18-Tc carrying the G to T and CGG to GCC in the *natT* allele as *Hind*III-*Xba*I fragment | Point mutation inserted into pEX18-Tc-*natT*$_{E29D}$ with primers L and M | This study |
| pEX18-Tc-Δ*natT* | pEX18-Tc carrying *natT* (PA1030) deletion cassette as *Hind*III-*Kpn*I fragment, Tet$^{R}$ | Deletion cassette generated by ligation of upstream fragment (primers N and O, *Hind*III-*Xba*I) and downstream fragment (primers P and Q, *Xba*I-*Kpn*I) amplified from PAO1 gDNA | This study |
| pEX18-Tc-Δ*natR* | pEX18-Tc carrying *natR* (PA1029, full length corrected gene) deletion cassette as *Hind*III-*Kpn*I fragment, Tet$^{R}$ | Deletion cassette generated by ligation of upstream fragment (primers T and U, *Hind*III-*Xba*I) and downstream fragment (primers R and S, *Xba*I-*Kpn*I) amplified from PAO1 gDNA | This study |
| pEX18-Tc-Δ*natRT* | pEX18-Tc carrying *natR* (PA1029, full length corrected gene) and natT (PA1030) deletion cassette as *Hind*III-*Kpn*I fragment, Tet$^{R}$ | Deletion cassette generated by ligation of upstream fragment (primers T and U, *Hind*III-*Xba*I) and downstream fragment (primers P and Q, *Xba*I-*Kpn*I) amplified from PAO1 gDNA | This study |
| pEX18-Tc-Δ*relA* | pEX18-Tc carrying *relA* (PA0934) deletion cassette as *Hind*III-*Kpn*I fragment, Tet$^{R}$ | Deletion cassette generated by ligation of upstream fragment (primers V and W, *Hind*III-*Xba*I) and downstream fragment (primers X and Y, *Xba*I-*Kpn*I) amplified from PAO1 gDNA | This study |

| | | | |
|---|---|---|---|
| pEX18-Tc-Δ*spoT* | pEX18-Tc carrying *spoT* (PA5338) deletion cassette as *Hin*dIII-*Kpn*I fragment, $Tet^R$ | Deletion cassette generated by ligation of upstream fragment (primers Z and AB, *Hin*dIII-*Xba*I) and downstream fragment (primers AC and AD, *Xba*I-*Kpn*I) amplified from PAO1 gDNA | This study |
| pEX18-Tc-Δ*nrtR* | pEX18-Tc carrying *nrtR* (PA4916) deletion cassette as *Hin*dIII-*EcoR*I fragment, $Tet^R$ | Deletion cassette generated by ligation of upstream fragment (primers AF and AG, *Hin*dIII-*Xba*I) and downstream fragment (primers AH and AI, *Xba*I-*EcoR*I) amplified from PAO1 gDNA | This study |
| pEX18-Tc-Δ*nadD2* | pEX18-Tc carrying *nadD2* (PA4917) deletion cassette as *Hin*dIII-*Kpn*I fragment, $Tet^R$ | Deletion cassette generated by ligation of upstream fragment (primers AJ and AK, *Hin*dIII-*Xba*I) and downstream fragment (primers AL and AM, *Xba*I-*Kpn*I) amplified from PAO1 gDNA | This study |
| pEX18-Tc-*pcnB1::gfp* | pEX18-Tc carrying construct for chromosomal insertion the *gfp* gene with its own Shine-Dalgarno downstream *pcnB1* (PA4919) gene as *BamH*I-*Kpn*I fragment, $Tet^R$ | Fragment generated by ligation of *gfp* containing fragment (from miniCTX::*pX2-gfp* as *BamH*I-*Kpn*I ) and upstream fragment (primers AN and AO, *Hind*III-*BamH*I) and downstream fragment (primers AP and AQ, *SphH*I-*Kpn*I) amplified from PAO1 gDNA | This study |
| pEX18-Tc-*natRT::gfp* | pEX18-Tc carrying construct for chromosomal insertion the *gfp* gene with its own Shine-Dalgarno downstream *natT* gene as *BamH*I-*Kpn*I fragment, $Tet^R$ | Fragment generated by ligation of *gfp* containing fragment (from miniCTX::*pX2-gfp* as *BamH*I-*Kpn*I ) and upstream fragment (primers AR and AS, *Hind*III-*BamH*I) and downstream fragment (primers N and AT, *Xho*I-*Kpn*I) amplified from PAO1 gDNA | This study |
| pEX18-Tc-Δ*natT::gfp* | pEX18-Tc carrying *natT* (PA1030) deletion cassette as *Hin*dIII-*Kpn*I fragment for deletion of *natT* in $natRT_{E29D}$*::gfp* strain, $Tet^R$ | Deletion cassette generated by ligation of upstream fragment (primers AJ and AK, *Hin*dIII-*Xba*I) and downstream fragment (primers AV and AU, *Xba*I-*Kpn*I) amplified from *natRT::gfp* gDNA | This study |
| pEX18-Tc- *natT-FLAG* | pEX18-Tc carrying construct for chromosomal insertion of 3xFLAG-tag at the C- terminus of NatT as *BamH*I-*Kpn*I fragment, $Tet^R$ | Fragment generated by ligation of upstream fragment (primers AR and AW, *Hin*dIII-*Xba*I) and downstream fragment (primers P and Q, *Xba*I-*Kpn*I) amplified from *natRT::gfp* gDNA | This study |

**Supplementary Table S4:** Oligos

| **Primers** | **Description** | **Sequence** |
|---|---|---|
| A | NatT_over_F | cca**GAATTC**GTGAGCGAGATCTGGCGACAGTGCAA |
| B | NatT_over_R | cc**GGTACC**TCACGCCGGATGCGGCAACTTTCCGT |
| C | NatT$_{ATG}$_over_F | cca**GAATTC**ATGAGCGAGATCTGGCGACAGTGCAA |
| D | NatR_over_F | cca**GAATTC**ATGAGCGCTCTCATCAAGGAACGTCC |
| E | NatR_over_R | cc**GGTACC**TCAGACCTTGCCGCGGATCGCATCGA |
| F | NatT_FLAG_R | cc**GGTACCTCACTTGTCATCGTCATCCTTGTAATCGA**<br>**TATCATGATCTTTATAATCACCGTCATGGTCTTTGTAGTC**C<br>GCCGGATGCGGCAACTTTCCGTCCA |
| G | pQF_natR_F | gat**AAGCTT**ATGAGCGCTCTCATCAAGGAACGTC |
| H | pQF_natR_R | gcg**GGATCC**TCAGACCTTGCCGCGGATCGCATC |
| I | RSF_His30F | gcg**GGATCC**GAGCGAGATCTGGCGACAGTGCAAGGG |
| J | RSF_His30R | cca**GAATTC**TCACGCCGGATGCGGCAACTTTCCGT |
| K | RSF_29F | cga**GATATC**GAGCGCTCTCATCAAGGAACGTCCCAG |
| L | NatT_R82A_F | CTTGCGCTGGGGGTCG**gcc**TTTGGCCGTCGCC |
| M | NatT_R82A_F | AAA**ggc**CGACCCCCAGCGCAAGGGCGGATAACG |
| N | NatT_KO_UPF | gat**AAGCTT**CTCTCTATCACGCCGGTCAATGCTTG |
| O | NatT_KO_UPR | cg**TCTAGA**CTTCCGCCCCTCGCTTCCTGTGGTGT |
| P | NatRT_KO_DnF | cg**TCTAGA**GTGGAGGGGGCTCGGTTGCCAGCGAT |
| Q | NatRT_KO_DnR | gc**GGTACC**GATGGAGTGGGGCTATGGGACAGATT |
| R | NatR_KO_DnF | cg**TCTAGA**TACCCGGCGCGCCTGTACGGGGGCCT |
| S | NatR_KO_DnR | gc**GGTACC**CGCTGCGCATGGCACTTCCCAGTTCC |
| T | NatRT_KO_UPF | gat**AAGCTT**AGGGTTCCTGGGATACCAGGACGGTG |
| U | NatRT_KO_UPR | cg**TCTAGA**GACGACCTCCTCATGTGTGTTTACA |
| V | relA_ko_UPF | gat**AAGCTT**ACTGGCAACTGACTCTGGCATA |
| W | relA_ko_UPR | cg**TCTAGA**CATCTTGCCTACCCTTTACCAC |
| X | relA_ko_DnF | cg**TCTAGA**TGAGGCGAGGCGGAAACAGGCC |
| Y | relA_ko_DnR | gc**GGTACC**TTCGTCCAGCCCCTCGCGAACC |
| Z | spoT_KO_UPF | gat**AAGCTT**GCCCAGCAGGTGCGCCGGCTGA |
| AB | spoT_KO_UPR | cg**TCTAGA**CAAGGGTTCACCCCCTGCCCGT |
| AC | spoT_KO_DnF | cg**TCTAGA**TGACCCGCTTTTTCCTGTGTCA |
| AD | spoT_KO_DnR | gc**GGTACC**CAGTTCGAAGCCGTTGCTGAAG |
| AF | nrtR_KO_UPF | gat**AAGCTT**CATGAATCGGTGATCCGCCGGGCGC |
| AG | nrtR_KO_UPR | cg**TCTAGA**CATCGTCACTCCTCTCTTCAGACCG |
| AH | nrtR_KO_DnF | cg**TCTAGA**TGACAAACGGGTGCGGCAGGGCCGG |
| AI | nrtR_KO_DnR | cca**GAATTC**TGCCGCGATAGAAGCCGCACGGGCG |
| AJ | PA4917_KO_UPF | gat**AAGCTT**CACATGGCTCACGCCATTCTGCCTC |
| AK | PA4917_KO_UPR | cg**TCTAGA**AGAAAATACCTTCCACTGCGAAATTTTG |
| AL | PA4917_KO_DnF | cg**TCTAGA**AGAGAGGAGTGACGATGAGTTCAGC |
| AM | PA4917_KO_DnR | gc**GGTACC**CTGGACTTCCTGGCCGATGGCGCAC |
| AN | Pa4919GFP_UPF | gat**AAGCTT**CCGCACCCAGGAGGAAGTGGTGCAT |
| AO | Pa4919GFP_UPR | gcg**GGATCC**TTAGGCCGGCACGTTGAAGACGTGG |
| AP | Pa4919GFP_DnF | cc**GCATGC**TCCTTTCGCACCTAACGGGAGCCCG |

| | | |
|---|---|---|
| AQ | Pa4919GFP_DnR | cc**GGTACC**ATGCTTGGCCAGGCCGCTGAGCGGA |
| AR | 10458_Pa1030GFP_UPF | gat**AAGCTT**AACAGGCGTTGCTCGAGGAACTGCTC |
| AS | 10459_Pa1030GFP_UPR | gcg**GGATCC**TCACGCCGGATGCGGCAACTTTCCGT |
| AT | 10602_Pa1030GFP_DnF | cca**CTCGAG**GTGGAGGGGGCTCGGTTGCCAGCGAT |
| AU | 18532_30KO_DnF | cg**TCTAGA**GGATCCTCTAGATTTAAGAAGGAGATA |
| AV | 18533_30KO_DnR | gc**GGTACC**AGGGCAGATTGTGTGGACAGGTAATG |
| AW | PA1030_C-FLAG-R | cg**TCTAGA**TCACTTGTCATCGTCATCCTTGT |

**Supplementary Table S5:** Crystallographic data collection and refinement statistics.

| | NatRT wt (PDB ID: 8QNL) | NatRT_E29D (PDB ID: 8QNQ) |
|---|---|---|
| Resolution range | 53.4 - 2.2 (2.3 - 2.2) | 53.7 - 2.3 (2.4 - 2.3) |
| Space group | P 1 21 1 | P 1 21 1 |
| Unit cell | 55.1 106.9 93.49 90 104.2 90 | 55.4 107.7 93.5 90 104.0 90 |
| Total reflections | 330825 | 269486 |
| Unique reflections | 47668 (4791) | 39154 (4174) |
| Multiplicity | 6.9 | 6.9 |
| Completeness (%) | 98.88 (99.50) | 92.23 (98.93) |
| Mean I/sigma(I) | 13.0 | 11.8 |
| Wilson B-factor | 43.19 | 43.74 |
| R-merge | 0.09 | 0.225 |
| R-pim | 0.04 | 0.094 |
| CC1/2 | 0.99 | 0.97 |
| Reflections used in refinement | 47663 (4791) | 39148 (4174) |
| Reflections used for R-free | 2322 (254) | 1971 (215) |
| R-work | 0.20 (0.27) | 0.23 (0.31) |
| R-free | 0.25 (0.34) | 0.26 (0.35) |
| Number of non-hydrogen atoms | 7857 | 7812 |
| macromolecules | 7708 | 7730 |
| ligands | 20 | 20 |
| solvent | 129 | 62 |

| | | |
|---|---|---|
| **Protein residues** | 980 | 985 |
| **RMS(bonds)** | 0.003 | 0.004 |
| **RMS(angles)** | 0.59 | 0.69 |
| **Ramachandran favored (%)** | 98.04 | 96.51 |
| **Ramachandran allowed (%)** | 1.76 | 2.88 |
| **Ramachandran outliers (%)** | 0.21 | 0.62 |
| **Rotamer outliers (%)** | 2.00 | 0.12 |
| **Clashscore** | 4.18 | 7.59 |
| **Average B-factor** | 52.07 | 51.72 |
| **macromolecules** | 52.23 | 51.85 |
| **ligands** | 42.38 | 38.85 |
| **solvent** | 43.91 | 39.79 |

Statistics for the highest-resolution shell are shown in parentheses.
